# Cell-Cell Adhesion During Nephron Development Is Driven by Wnt/PCP Formin Daam1

**DOI:** 10.1101/2020.08.18.256123

**Authors:** Vanja Krneta-Stankic, Mark Corkins, Adriana Paulucci-Holthauzen, Malgorzata Kloc, Andrew Gladden, Rachel Miller

## Abstract

E-cadherin junctions facilitate the assembly and disassembly of cell-cell contacts that drive development and homeostasis of epithelial tissues. The stability of E-cadherin-based junctions highly depends on their attachment to the actin cytoskeleton, but little is known about how the assembly of junctional actin filaments is regulated. Formins are a conserved group of proteins responsible for the formation and elongation of filamentous actin (F-actin). In this study, using *Xenopus* embryonic kidney and Madin-Darby canine kidney (MDCK) cells, we investigate the role of the Wnt/ planar cell polarity (PCP) formin protein Daam1 (Dishevelled-associated activator of morphogenesis 1) in regulating E-cadherin based intercellular adhesion. Using live imaging we show that Daam1 localizes to newly formed cell-cell contacts in the developing nephron. Furthermore, analyses of junctional F-actin upon Daam1 depletion indicate a decrease in microfilament localization and their slowed turnover. We also show that Daam1 is necessary for efficient and timely localization of junctional E-cadherin, which is mediated by Daam1’s formin homology domain 2 (FH2). Finally, we establish that Daam1 signaling is essential for promoting organized movement of renal cells. This study demonstrates that Daam1 formin junctional activity is critical for epithelial tissue organization.

## INTRODUCTION

Extensive cellular rearrangements with changes in cell shape drive morphogenesis, including for example, the process of tubulogenesis. To execute these processes successfully, cells must be able to interact with each other and their environment in a timely and coordinated manner. These interactions entail transduction of specific signals arising from adhesive contacts with the extracellular matrix (ECM) and neighboring cells. How cells remodel their adhesions is one of the central questions in epithelial tissue biology.

The cadherin family of cell adhesion proteins such as E-cadherin facilitate intercellular adhesion and formation of cellular junctions (Adams et al., 1998; Takeichi, 2014; Yap et al., 2015). Changes in E-cadherin-based adhesion are associated with developmental disorders and progression of disease (Friedl and Mayor, 2017; Mendonsa et al., 2018). E-cadherin levels at intercellular contacts depend on the organization of the actin cytoskeleton, but much remains to be learned about how the actin assembly is regulated at these adhesive sites. Moreover, much of our understanding of *in vivo* actin regulation and dynamics in cell-cell adhesion derives from observations in cell culture systems, invertebrate embryos and the vertebrate skin. Understanding of junction dynamics in intact vertebrae tissues is challenging, due to technical limitations and tissue inaccessibility. In this study, we probe the role of the formin protein Dishevelled-associated activator of morphogenesis 1 (Daam1) in intercellular adhesion during kidney development using *Xenopus laevis* embryonic kidney and Madin-Darby canine kidney (MDCK) cells.

Similar to many organs in our body, the kidney consists of a network of epithelial tubules. The epithelial tubules of the kidney are called nephrons whose morphology is vital to kidney function. Mesenchymal-epithelial transitions (MET) and coordinated cell rearrangements facilitate nephron morphogenesis. Nephrons arise from the mass of mesenchymal cells that undergo MET to form tubules consisting of tightly connected epithelial cells (McMahon, 2016; Saxen, 1987). Oriented cell intercalations drive the elongation of nephric tubules through a process called convergent-extension (CE) (Castelli et al., 2013; Karner et al., 2009; Kunimoto et al., 2017; Lienkamp et al., 2012). Extensive cytoskeletal rearrangements characterized by changes in cell shape and coordinated cell movements accompany CE. Although E-cadherin-based adhesions are implicated in mediating both MET and maintenance of coordinated cell rearrangements (Campbell and Casanova, 2016), very little is known about how they function in nephrogenesis (Combes et al., 2015; Lefevre et al., 2017; Marciano et al., 2011; Vestweber et al., 1985).

Daam1 is a formin protein required for nephric tubulogenesis (Miller et al., 2011). Formin proteins coordinate the organization of the actin cytoskeleton by nucleating and polymerizing unbranched actin filaments. While Rho GTPases activate most formins, the activation of Daam1 depends on its interaction with Dishevelled (Dvl), a key intracellular component of the Wnt signaling pathway (Liu et al., 2008). The Wnt signaling pathway plays important roles in nephron development (McMahon, 2016; Miller and McCrea, 2009). The secreted Wnt ligands bind Frizzled (Fz) receptors and subsequently, via Dvl regulate the canonical (β--catenin-dependent) and non-canonical (β--catenin-independent)/planar cell polarity (PCP) signaling. While the canonical signaling commonly governs inductive events and cell fate, the non-canonical/PCP branch is associated with influencing cell behaviors and morphology. Nonetheless, the roles for different branches of the Wnt pathway continue to evolve as recent studies provide evidence for cross-talk between these two branches (Nagy et al., 2016; O’Brien et al., 2018). Dvl regulates the non-canonical/PCP branch of the Wnt pathway through direct interaction with Daam1 (Liu et al., 2008).

Increasing evidence suggests that formins may function as key regulators of the actin assembly at the cell-cell junctions (Grikscheit and Grosse, 2016). Recent work in a mouse mammary gland epithelial cell line, for example, has indicated that Daam1 is important for the stability of epithelial cell junctions (Nishimura et al., 2016). Here, we expand on these findings by examining the functional role of Daam1 in cellular junctions in the context of tissue morphogenesis by analyzing its role in nephron development. Furthermore, using live cell imaging we show that during establishment of cellular junctions, Daam1 first localizes to cellular protrusions that initiate cell-cell contact, and subsequently, to newly formed junctions to promote their stability. We find that Daam1 facilitates nephron morphogenesis by regulating the assembly of junctional filamentous actin (F-actin) and in turn promotes the E-cadherin-based epithelial adhesion.

## RESULTS

### Daam1 co-localizes with F-actin and E-cadherin within the nephric primordium

Knockdown of Daam1 in *Xenopus* disrupts nephron morphology without apparent effect on the expression of genes related to early differentiation events (Miller et al., 2011). Differentiation signals driving development of the nephric mesoderm in *Xenopus* largely function before the onset of tubular morphogenesis (Vize et al., 2003). To further probe the mechanism by which Daam1 regulates the shaping of nephrons, we analyzed the subcellular localization of fluorescently tagged Daam1 at the beginning of tubular morphogenesis (around NF stage 30). Through kidney-targeted microinjections that were targeted to early embryonic cells fated to contributed to kidney (presumptive nephron progenitors) (DeLay et al., 2016; Moody and Kline, 1990), we expressed 1ng of fluorescently tagged Daam1 mRNA in the presumptive nephron progenitors and analyzed its localization in fixed and live tissue.

For analyses of fixed tissue, embryos were subjected to whole-mount staining. The samples were stained with an antibody against GFP to visualize Daam1, and an antibody against Lhx1 to label nephron progenitors (DeLay et al., 2018; Venegas-FERRÍN et al., 2010). An additional staining with Phalloidin allowed us to visualize the F-actin cytoskeleton, or alternatively, another antibody was used to define the localization of E-cadherin **(Figure 1)**. Consistent with its previously reported subcellular localization in other cell types (Corkins et al., 2019; Higashi et al., 2019; Jaiswal et al., 2013; Kawabata Galbraith et al., 2018; Kida et al., 2007; Nishimura et al., 2012, 2016), Daam1 co-localizes with patches of F-actin, showing a more diffuse staining pattern in the cytoplasm and a strong localization to cell junctions **(Figure 1A)**. Furthermore, co-immunostaining for GFP and E-cadherin demonstrated that E-cadherin is expressed within the nephric primordium and it co-localizes with Daam1 at the cell-cell junctions **(Figure 1B)**.

**Figure 1.**
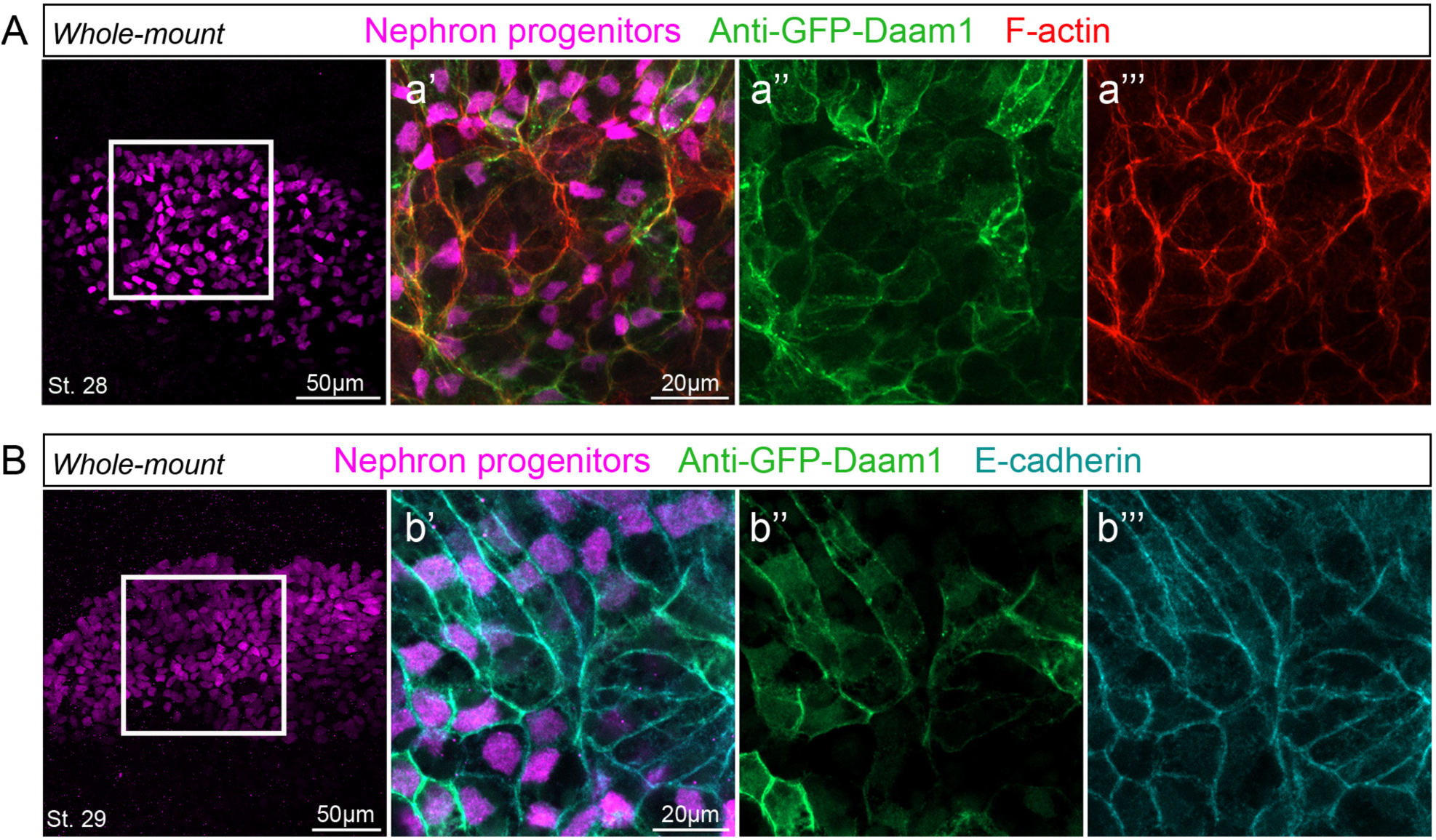
Daam1 co-localizes with junctional F-actin and E-cadherin during early nephron development. Confocal maximum image projections of whole-mount immunostaining of *Xenopus* nephric primordium labeled by Lhx1(magenta) and GFP to visualize Daam1 (green) in conjunction with, (A) Phalloidin staining to visualize F-actin (red) or (B) E-cadherin (cyan); a’-a’’’ and b’-b’’’ represent close-up images of white boxes.

During the early stages of development, the opaqueness of the *Xenopus* epithelium hinders imaging of fluorescent protein expression in internal tissues, including the pronephric primordium. In fixed tissues, this is overcome by using clearing agents such as BA:BB (1:2 mixture of benzyl alcohol and benzyl benzoate, aka Murray’s clear). However, this process requires dehydration of samples in methanol or ethanol prior to clearing, which is incompatible with the use of fluorescent phalloidins (Becker and Gard, 2006). To overcome this obstacle, we briefly washed embryos (< 20 sec) in isopropanol prior to clearing with BA:BA (Nworu et al., 2014; Strickland et al., 2004). This allowed us to visualize Daam1 in conjunction with F-actin in intact pronephric primordium **(Figure 1A)**.

To overcome an analogous challenge in live embryos, we developed a novel way of imaging the nephric primordium *in vivo* **(Figure 2A)**. Previous studies have used the “windowed” embryo approach consisting of microsurgical removal of the surface ectoderm to expose and image underlying tissue (Kim and Davidson, 2013). Adopting this approach, we created “kidney-windowed” embryos by removing the surface epithelium and exposing the underlying nephric primordium for high-resolution *in vivo* imaging. *In vivo* time-lapse imaging of “kidney-windowed” embryos showed GFP-Daam1 localizing to the cell-cell junctions. However, we observed that GFP-Daam1 also localized to cytoplasmic vesicles and cellular protrusions **(Figure 2B, Video S1)**. These observations are in line with previous reports on Daam1’s localization and likely hindered in imaging of fixed tissue due to unfavorable fixation conditions for observations of cytoplasmic vesicles and cellular protrusions (Corkins et al., 2019; Jaiswal et al., 2013; Kawabata Galbraith et al., 2018; Kida et al., 2007; Nishimura et al., 2012, 2016). Moreover, to better understand the dynamics of Daam1 in the context of cell junctions, we imaged *de novo* formation of cell-cell junctions in dissociated GFP-Daam1 expressing cells derived from the nephric primordia **(Figure 2A)**. *In vivo* time-lapse analyses of these cells showed that Daam1 localizes to filopodia-like protrusions and subsequently, to newly formed junctions **(Figure 2B, Video S2)**.

**Figure 2.**
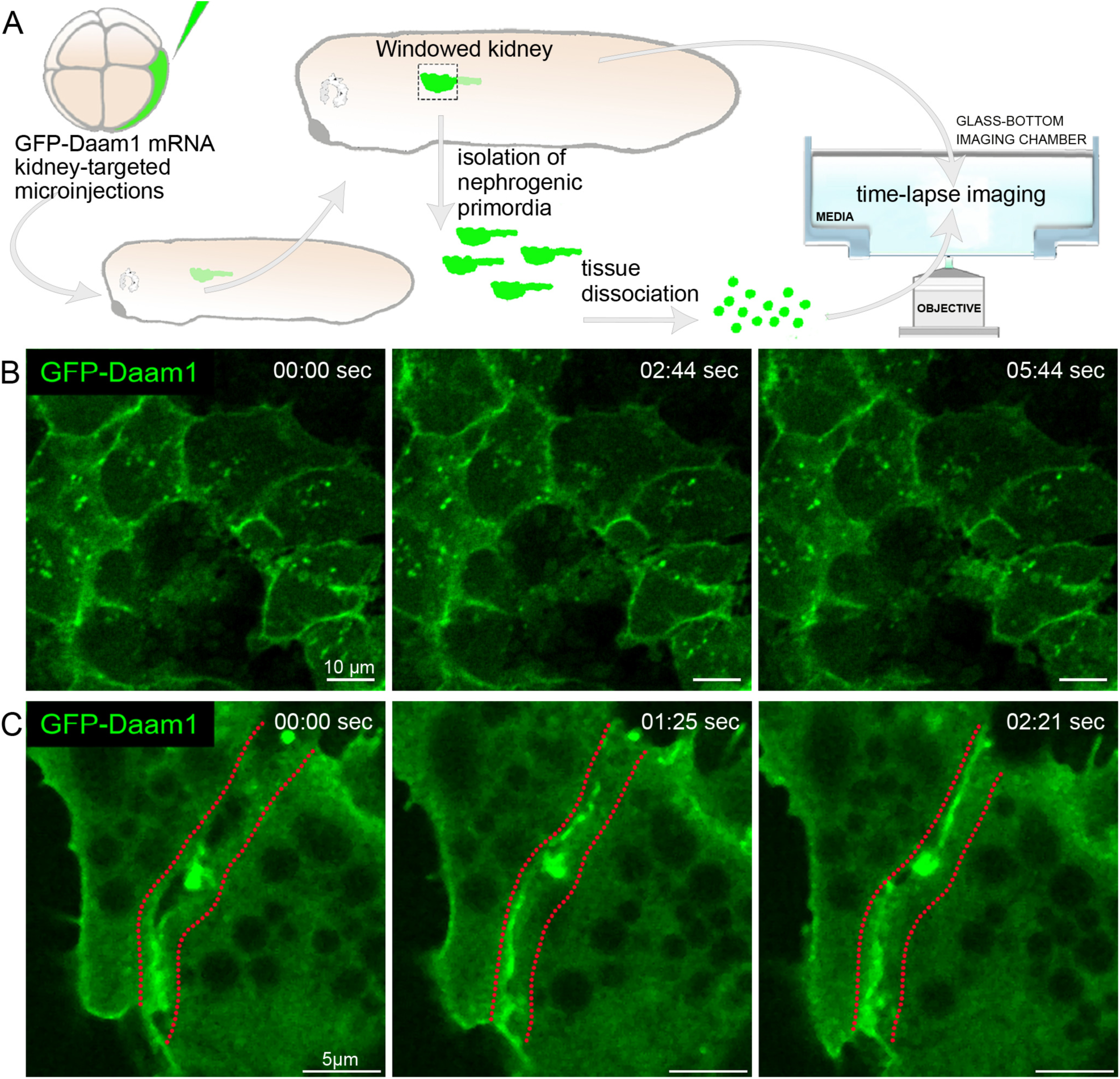
Daam1 localizes to newly formed cell-cell contacts. (A) Schematic illustration showing steps involved in preparation of “windowed kidney” embryos and primary cultures expressing GFP-Daam1. Please note that for clarity of illustration the 8-cell GFP-Daam1 injected *Xenopus* blastomere is fate-mapped strictly to the nephric primordium and that blastomere also contributes to epidermis, ventral and dorsal somites, hidgut, proctodeum and trunk neural crest cells (DeLay et al., 2016; Moody and Kline, 1990). (B) Time-lapse imaging montage of the nephric primordium expressing GFP-Daam1 in “windowed kidney” embryos. Elapsed time is indicated at the top in seconds; see **Video S1**. (C) Time-lapse imaging montage shows cells isolated from a developing nephron expressing GFP-Daam1 mRNA adhering with each other. Elapsed time is indicated at the top in seconds. The border arising between two cells is delineated by the red dotted line; see **Video S2**.

Finally, we assessed the localization of Daam1 in the mature epithelium of fully developed nephrons **(Figure S1)**. Interestingly and importantly, junctional localization of Daam1 was not detected in the mature epithelium. Taken together, these data suggested the potential role for Daam1 in regulating the intercellular adhesion of renal progenitors specifically at the onset of tubulogenesis and CE.

### Daam1 controls the organization and assembly of junctional F-actin within the nephric primordium

The observation that Daam1 co-localizes with junctional F-actin in developing *Xenopus* nephron, led us to ask whether Daam1 regulates F-actin. To address this, we depleted Daam1 in nephron progenitors by utilizing Morpholino (MO) oligos in kidney-targeted microinjections (DeLay et al., 2016; Moody and Kline, 1990). A proven Daam1 MO or an established Control (standard) MO (Habas et al., 2001; Liu et al., 2008; Miller et al., 2011) was co-injected with a membrane tagged GFP (mGFP) mRNA, that served as a linage tracer. MO-injected embryos were fixed at the onset of tubular morphogenesis (around NF stage 30) and subjected to whole-mount fluorescent staining and high-resolution confocal imaging. To verify the success of our injections and knockdowns, we also carried out Western blot analyses of total protein lysates prepared from stage 30 MO-injected embryos **(Figure 3B)**. Phalloidin staining revealed that upon Daam1 depletion, the F-actin in renal progenitors becomes disorganized and significantly reduced at cell-cell junctions **(Figure 3A and 3C, Videos S3 and S4)**. These results suggest that Daam1 contributes to the organization of the F-actin cytoskeleton and actin filaments at cell-cell junctions during nephron development.

**Figure 3.**
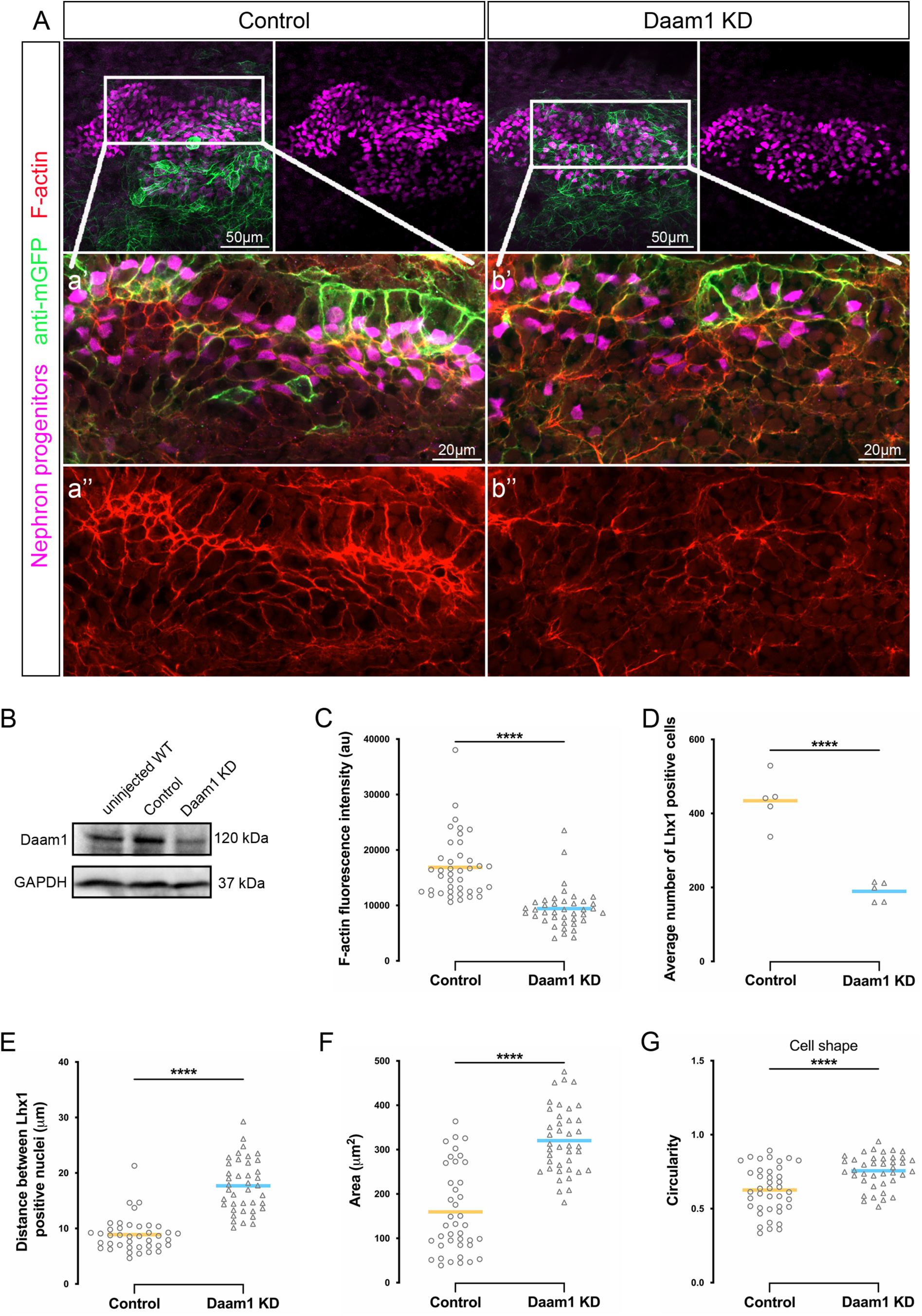
Effects of Daam1 depletion on the nephrogenic primordium. (A) Maximum projection confocal images of F-actin expression (red) in nephric primordium (magenta) in Control and Daam1 knockdown embryos. a-a’’ and b’-b’’ represent close-up images of the corresponding regions in white boxes; see **Videos S3 and S4**. (B) Western blot showing Daam1 and GAPDH (control) protein levels for uninjected wild type (WT) and Control (Standard morpholino) and Daam1 KD (Daam1 morpholino) injected embryos. (C) The graph showing the relative fluorescence intensity levels of junctional F-actin in the nephric primordia of Control and Daam1 KD embryos. N_Control_=40 junctions on 2 embryos and N_Daam1KD_=40 junctions on 2 embryos. ****P < 0.0001 analyzed by unpaired t-test. (D-G) Morphometric analyses of Control and Daam1-depleted nephric primordia. The thick bars represent the mean, ****P < 0.0001 analyzed by unpaired t-test, (E-G) N_Control_=40 junctions on 2 embryos and N_Daam1 KD_=40 junctions on 2 embryos. Graphs showing comparison between Control and Daam1-depleted nephric primordia of, (D) the average number of Lhx1-positive nephron progenitors where N_Control_=5 embryos and N_Daam1KD_=5 embryos, (E) the relative distance between nearest neighbors of Lhx1-positive nuclei, (F) the relative cell area and (G) the relative circularity, where 1 represents the perfect circle.

We also observed alterations in spatial positioning and the overall organization of nephron progenitors upon depletion of Daam1. Nephric primordia in Daam1-morphants consist of fewer progenitor cells **(Figure 3D)** that are spaced farther part from one another **(Figure 3E)** compared to control animals. In addition to changes in the number and position of nephron progenitors, we also noted changes in cell morphology. Nephric progenitors with diminished Daam1 activity display an increase in cell area **(Figure 3F)** and circularity **(Figure 3G)**.

Since polymerization of F-actin filaments is a highly dynamic process, and to better understand possible mechanisms underlying the observed morphological perturbations, we next assessed the dynamic behavior of F-actin *in vivo* using fluorescence recovery after photobleaching (FRAP) assays. We labeled F-actin in developing nephrons by co-injecting mCherry tagged Utrophin (mCherry-UtrCH) mRNA (Burkel et al., 2007) along with control or Daam1 morpholino. To probe F-actin dynamics in the context of the intact animal, we utilized the “kidney-windowed” approach in NF stage 30 embryos. Overall, FRAP experiments suggest slower turnover of junctional F-actin in Daam1-depleted nephrons in comparison with controls **(Figure 4)**. In control nephrons, 100% of bleached cell junctions successfully recover fluorescence signal. Whereas, in Daam1-depleted nephrons, only 59% of bleached junctions recovered **(Figure 4A)**. Cell-cell junctions are comprised of dynamically mosaic E-cadherin clusters coupled to different actin dynamics (Cavey et al., 2008; Indra et al., 2018). Therefore, the differences observed in the dynamics of F-actin in Daam1 knockdown contexts potentially indicate the existence of two actin pools differentially regulated by Daam1. However, it also possible that the observed differences stem from the cell heterogeneity (e.g. in respect to Daam1 expression levels or cell-type representation) within the nephric primordium. As we were unable to quantitatively assess F-actin dynamics in junctions that fail to recover, we only used junctions with detectable recovery signal to determine the mean half-time to recovery **(Figure 4D)** and the mean mobile fraction **(Figure 4E)** values. For each junction values were taken from individually fitted curves. These data indicate that the mean recovery half-time for junctional F-actin in Daam1 KD nephrons is significantly slower (5.57 sec ± 0.99 s.e.m.) in comparison with control nephrons (3.70 sec ± 0.35 s.e.m.) **(Figure 4D)**. In contrast, the mean mobile fractions are relatively similar (56.0% ± 4.2 s.e.m for control and 54.3 % ± 4.5 s.e.m for Daam1 KD) **(Figure 4E)**. From these data, we conclude that Daam1 is driving the rate of F-actin turnover to promote polymerization of junctional actin during nephron development. Moreover, these data also suggest that the decrease in F-actin fluorescence levels observed upon Daam1 depletion is likely a consequence of impaired actin assembly.

**Figure 4.**
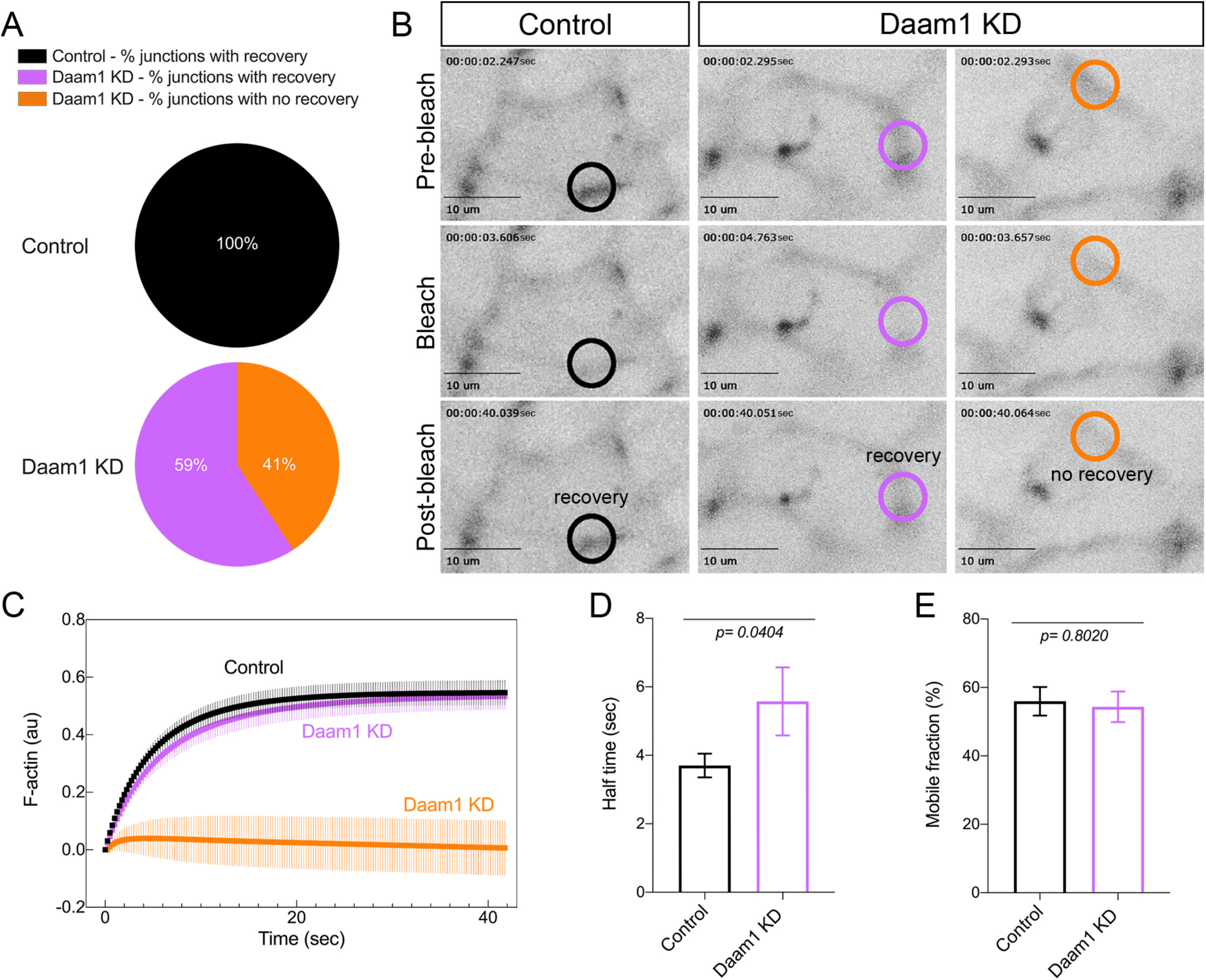
Daam1 regulates assembly of junctional F-actin in developing nephron. F-actin dynamics at cell-cell junctions of Control and Daam1-depleted developing nephrons expressing mCherry-Utrophin were assessed using FRAP. (A) Percentage of junctions showing recovery of fluorescence after bleaching in Control (black, N_total_=27 junctions, 1-5 junctions/embryo) and Daam1 KD (purple and orange, N_total_=27 junctions, 1-5 junctions/embryo) nephrons. (B) Typical time-lapse images of Control and Daam1-depleted cell junctions before and after photobleaching. In each image, the bleached region is highlighted with a circle (black - Control junction showing recovery, purple - Daam1 KD junction showing recovery and orange - Daam1 KD junction showing no-recovery of fluorescence after photobleaching). (C) Graph shows average recovery curves obtained from individual best-fit plots for Control (black), Daam1 KD junctions with (purple) and without (orange) recovery of fluorescence after photobleaching. (D-E) Bar graphs comparing Control and Daam1 KD profiles calculated from individual best-fit curves for Control (black) and Daam1 KD junctions with recovery of fluorescence after photobleaching (purple). Data represent the mean ± S.E. from three independent experiments. P-values were analyzed by unpaired t-test. (D) Bar graph of the relative half-times for F-actin. (E) Bar graph of the relative mobile fraction for F-actin.

### Daam1 promotes localization of E-cadherin at cell-cell contacts

The interplay between E-cadherin and the actin cytoskeleton promotes intercellular adhesion and assembly of cellular junctions. Therefore, we asked whether the changes observed in junctional F-actin dynamics upon the knockdown of Daam1 alters the intercellular adhesion between nephron progenitors. To examine if Daam1 regulates intercellular adhesion, we analyzed the effect of Daam1 depletion on E-cadherin localization in developing nephron **(Figure 5)**. Daam1 morphants displayed reduced levels of E-cadherin at the interfaces between neighboring cells during these early stages of pronephric morphogenesis **(Figure 5A)**. This difference was quantified and likewise made evident by measuring the fluorescence intensity profiles of E-cadherin along the length of individual junctions **(Figure 5B)**. Of note and in contrast, there was no difference in the overall E-cadherin protein levels between Daam1 knockdown and control embryos as determined by Western blotting **(Figure 5C)**. These findings suggest that Daam1 is likely more important for localization of E-cadherin at cell-cell contacts as opposed to regulating the overall expression levels of E-cadherin.

**Figure 5.**
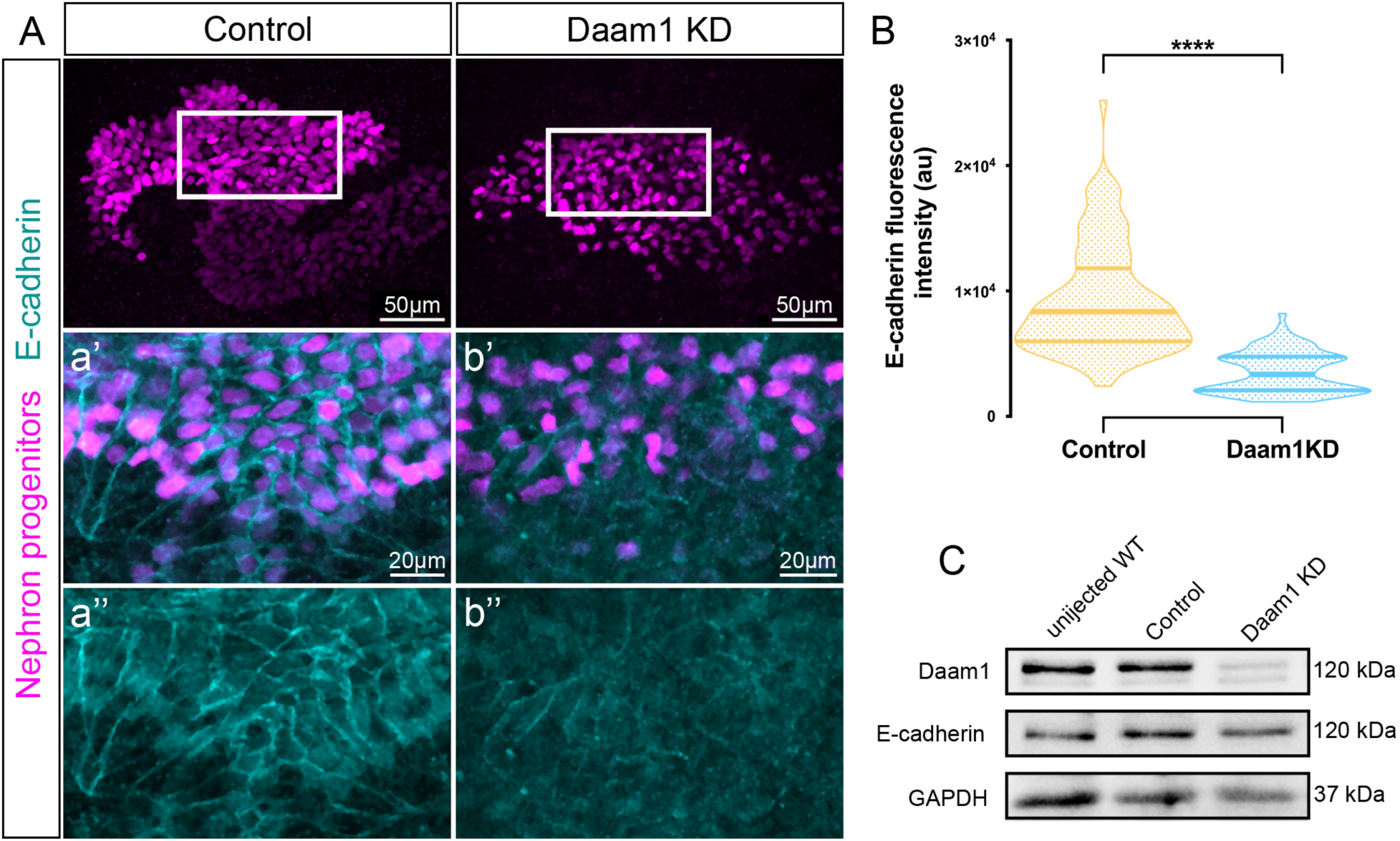
Daam1 promotes localization of junctional E-cadherin. (A) Maximum projection confocal images of E-cadherin expression (cyan) in nephric primordium (magenta) in Control and Daam1 KD embryos a-a’ and b-b’ represent the close-up images of corresponding regions in white boxes. (B) Violin plots depicting the relative fluorescence intensity of junctional E-cadherin in the nephric primordia of Control (orange) and Daam1 KD (blue). N_Control_=88 junctions on 4 embryos and N_Daam1 KD_=84 junctions on 4 embryos. Center-lines represents median; Limits show 1^st^ and 3^rd^ quartile. ^****^P<0.0001 analyzed by unpaired t-test. (C) Western blot showing Daam1, E-cadherin and GAPDH (control) protein levels in uninjected wild type (WT) and Control (Standard morpholino) and Daam1 KD (Daam1 morpholino) injected embryos.

Interestingly, Daam1 depleted cells remain capable of forming nephrons. Studies carried out in MDCK cells using the pan-formin inhibitor SMIFH2 suggest that formins are required for early, but not later stages of cell-cell adhesion (Collins et al., 2017). Moreover, Daam1 depleted mammary epithelial cells form a monolayer characterized by irregular tilting of lateral cell membranes and distorted cell morphology (Nishimura et al., 2016). To further understand the function of Daam1 in nephron assembly, we analyzed the epithelium of mature nephrons in NF stages 39-40 embryos **(Figure S2)**. Indeed, we did not observe any apparent changes in the local concentration of junctional E-cadherin in mature nephrons of Daam1 knockdown and control embryos **(Figure S2A)**. However, Daam1 knockdown nephrons displayed defects in the size of the tubular lumen. The diameter of tubular lumens was more variable in Daam1 deficient nephrons compared to the controls. Similar to observations in mammary epithelial cells (Nishimura et al., 2016), tubular cells in Daam1 depleted nephrons are less uniform in shape and characterized by an irregular tilting of lateral cell membranes **(Figure S2A; Videos S5 and S6)**. We further found that when visualized by transmission electron microscopy (TEM), Daam1 depleted cells appear less columnar, characterized by indistinct and wavy cell borders **(Figure S2B)**. Our results suggest that Daam1 regulates the adhesion between nephron progenitor cells and subsequently, the morphology of the mature nephric epithelium.

### E-cadherin localization is mediated by the Daam1 FH2 domain

Daam1 is known to act upstream of small Rho-GTPases, which regulate the actin cytoskeleton (Habas et al., 2001; Liu et al., 2008); therefore, we next tested the importance of the actin polymerization activity of Daam1 in intercellular adhesion. Formins are defined by a conserved Formin Homology 2 (FH2) domain. The Daam1 forms a dimer via its FH2 domain, responsible for nucleation and elongation of actin filaments (Lu et al., 2007; Yamashita et al., 2007). The mutation isoleucine-to-alanine (Ile698Ala) in the Daam1 FH2 domain abolishes the actin polymerization activity of Daam1 *in vitro* (Lu et al., 2007) and *in vivo* (Liu et al., 2008; Nishimura et al., 2016). In kidney targeted-injections, we expressed either full-length GFP-Daam1 or GFP-Daam1 FH2 mutant (Ile698Ala) mRNA and analyzed the effect on E-cadherin localization in the nephric progenitors **(Figure 6)**. Nephric progenitors expressing GFP-Daam1 FH2 mutant mRNA showed reduced levels of E-cadherin at the interfaces between neighboring cells in comparison to nephric progenitors expressing GFP-Daam1 **(Figure 6A-B)**. However, the E-cadherin phenotype appeared to be less prominent than in Daam1 knockdown. Functional studies showed that while an isoleucine-to-alanine mutation within FH2 domain abolishes Daam1’s ability to polymerize actin, it does not prevent its activation of Rho (Liu et al., 2008). This could be one possible explanation as to why E-cadherin localization is more affected in Daam1 knockdown nephric progenitors. Furthermore, we assessed Daam1 protein levels in injected embryos to determine if the observed differences in E-cadherin localization were potentially due to underlying disparities in translation efficiency or protein stability of GFP-Daam1 and GFP-Daam1 FH2 mutant **(Figure 6C)**. We found that Daam1 protein was present at equivalent levels in the two samples, making these possibilities unlikely. Ultimately, the GFP-Daam1 FH2 mutant expressing progenitors mature into nephrons characteristic of Daam1 knockdown **(Figure S3)**. These results establish that the Daam1 FH2 domain is necessary for the localization of E-cadherin to cell-cell contacts in nephron progenitors.

**Figure 6.**
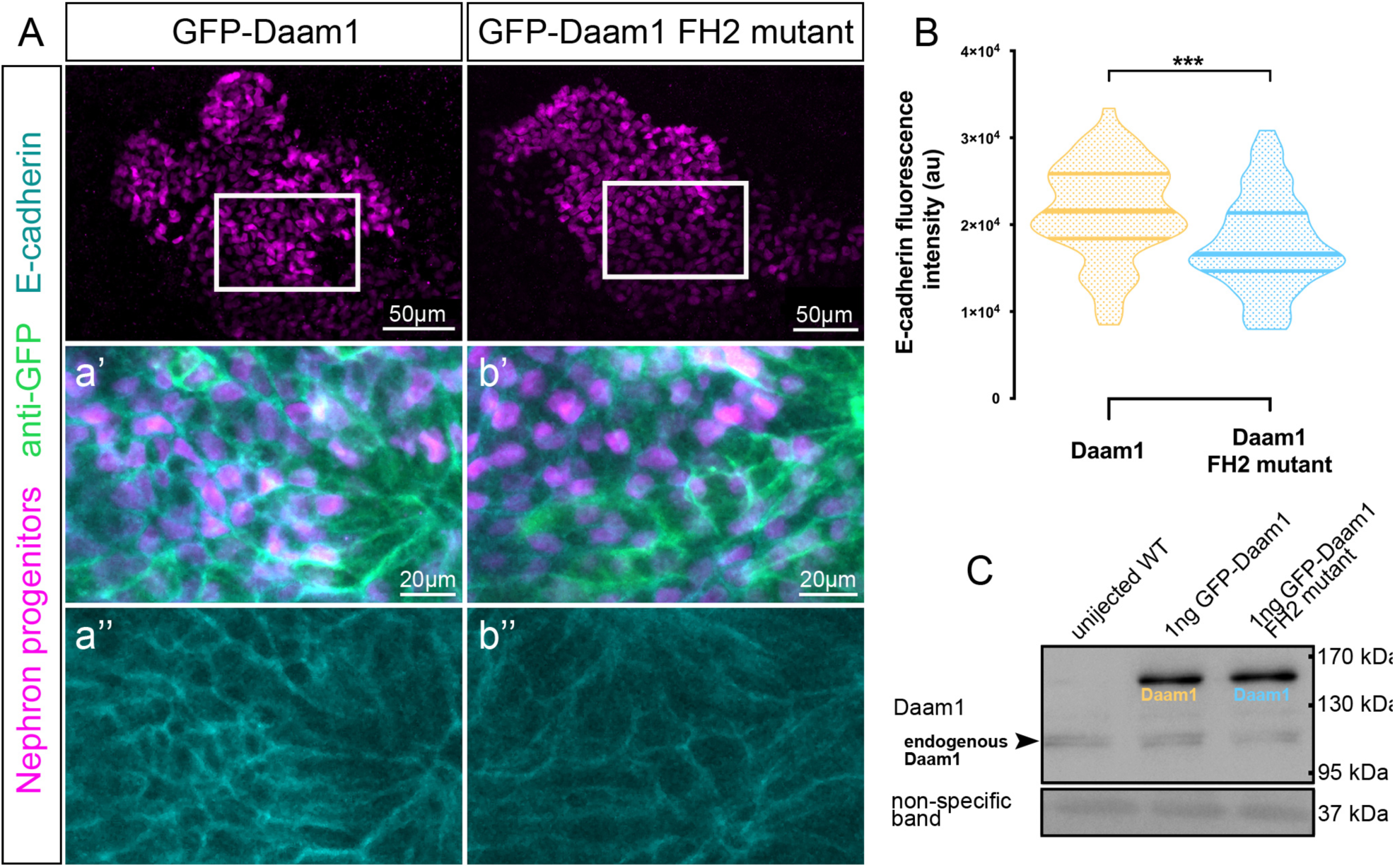
E-cadherin localization to cell-cell junctions is mediated by FH2 domain of Daam1. (A) Maximum projection confocal images showing E-cadherin staining (cyan) in nephric primordium (magenta) expressing GFP-Daam1 or GFP-Daam1 FH2 mutant mRNA. a-a’ and b-b’ represent close-up images of corresponding regions in white boxes. (B) Violin plots depicting the relative fluorescence intensity of junctional E-cadherin in the nephric primordia expressing GFP-Daam1 (orange) and GFP-Daam1 FH2 mutant (blue) mRNA. N_Daam1_=60 junctions on 3 embryos and N_Daam1FH2mutant_=55 junctions on 3 embryos. Center-lines represents median; Limits show 1^st^ and 3^rd^ quartile. ^***^P<0.0002 analyzed by unpaired t-test. (C) Western blot showing the exogenous and endogenous protein levels of Daam1 in uninjected wild type (WT) embryos, embryos injected with 1 ng of GFP-Daam1 mRNA and 1ng GFP-Daam1 FH2 mutant mRNA. The non-specific band confirms equal loading.

### Daam1 mediates cohesion of MDCK cells

Nephron morphogenesis is achieved through the process called convergent extension (CE) (Lienkamp et al., 2012). The CE is a type of collective cell movement characterized by a series of coordinated and directed cell rearrangements (Huebner and Wallingford, 2018; Tada and Heisenberg, 2012). In recent years E-cadherin has emerged as a key mediator for coordinating cohesion and directional persistence of collective cell movements in both epithelial and mesenchymal clusters (Cai et al., 2014; Campbell and Casanova, 2015; Cohen et al., 2016). Our results demonstrate that Daam1 is necessary for the organization of nephrogenic primordium **(Figure 3)** and a proper localization of E-cadherin **(Figures 5 and 6)**; therefore, we wanted to determine if Daam1 is necessary for the coordination of direction between renal cells. Because the opaqueness of *Xenopus* nephron progenitors prevents *in vivo* tracking of their movements in 3D, we utilized Madin-Darby Canine Kidney (MDCK) cells. We generated MDCK cells constitutively expressing an shRNA against Daam1 and analyzed whether the E-cadherin localization is affected in these cells **(Figures 7A and S4)**. The efficiency of shDaam1 knockdown was confirmed by Western blot **(Figure 7B)**. Similar to what we saw in *Xenopus* nephrons, we observed impaired localization of E-cadherin in nascent **(Figure 7A)**, but not mature adhesions **(Figure S4)** upon knockdown of Daam1. We next examined the migratory behavior of Daam1 knockdown cells in a time-lapse imaging of the wound-healing assay **(Figure 7C-F; Video S7)**. We found Daam1 knockdown cells exhibit a delay in a wound closure compared to control cells **(Figure 7C; Video S7)**. Additionally, we also observed random detachment of Daam1-deficient cells from the migrating epithelial sheets **(Video S7)**. To better understand the behavior of these cells, we tracked their movement within the migrating sheets over time. **(Figure S7)**. From these tracks, we obtained the relative distances over which cells traveled and used those distances to assess cell velocities **(Figure 7C)**. These analyses show that the speed at which Daam1 knockdown cells move is higher than that of the control cells, demonstrating that delayed wound closure in Daam1 knockdown cells is not caused by their slow movement. However, mapping the trajectory paths for Control **(Figure 7E)** and shDaam1 **(Figure 7D)** cells revealed that the movement of Daam1-deficient cells is less directed compared to control. These data demonstrate that Daam1 is necessary for communication of direction between the cells. Taken together, these results indicate that Daam1 contributes to cohesion by regulating connectiveness of cells through E-cadherin.

**Figure 7.**
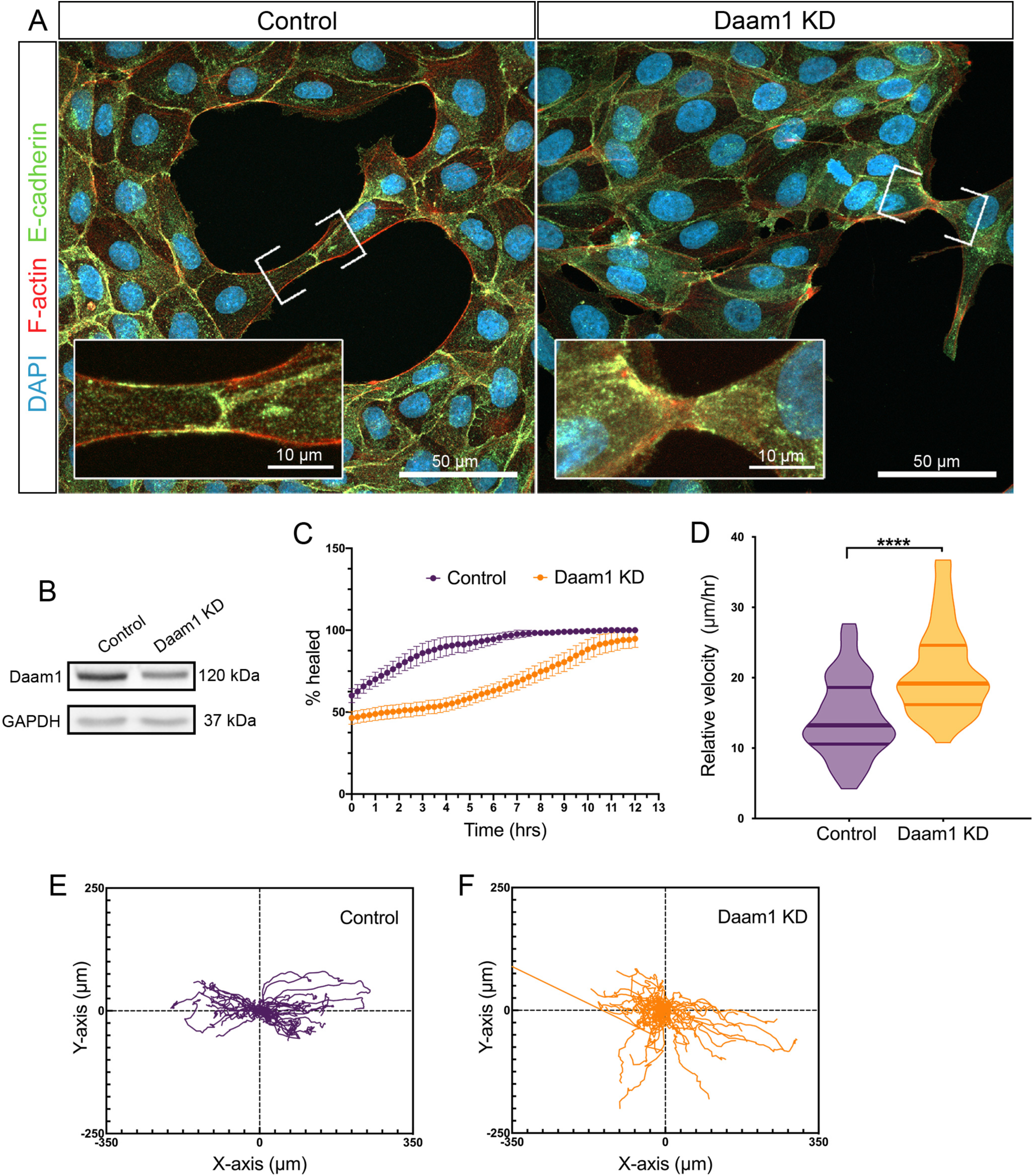
Daam1-depleted MDCK cells display compromised localization of E-cadherin at cell-cell contacts and impaired cohesion during collective movement. (A) E-cadherin (green), F-actin (red) and DAPI (blue) in subconfluent the MDCK Control and shDaam1 knockdown cells. E-cadherin localization in the nascent cell-cell contacts (marked by white brackets and shown enlarged in corresponding white boxes) is impaired in shDaam1-deficient cells. (B) Western blot analysis of Daam1 and GAPDH protein levels in the MDCK Control and shDaam1 knockdown cells. (C-F) Summary of the wound-healing experiments for the MDCK Control and Daam1 KD cells, **see Video S7**. (C) Daam1 depletion impairs wound closure. The graph represents the percent of the wound surface area over time for Control (purple) and Daam1 KD (orange) cells. Error bars indicate S.E. of the mean on 4 assays. (D-F) Manually tracking migration paths of single-cells during the wound closure demonstrates that Daam1 organizes collective movement of the MDCK epithelial monolayers by modulating the speed and directionality of individual cells. Depletion of Daam1 results in increased velocity and random migration. N_Control_=52 cells from 4 assays and N_Daam1 KD_=42 cells tracked from 4 assays. Cells were tracked in 15 minutes increments for 12 hours. (D) Violin plots represent migration velocity calculated from tracking traveled distances of single cells for Control and Daam1 KD cells. Center-lines represents median; Limits show 1^st^ and 3^rd^ quartile. ^***^P<0.0001 analyzed by unpaired t-test. (E) Wind-rose plot showing migration tracks of individual Control cells. (F) Wind-rose plot showing migration tracks of Daam1 KD cells.

## DISCUSSION

Using *Xenopus* embryonic kidney and MDCK cells as model systems, we show that Daam1 mediates E-cadherin dependent intercellular adhesion and organization of nephrogenic primordia by regulating polymerization of junctional actin filaments. Ultimately, this affects the morphology of mature nephric epithelium. These findings have a number of important implications for the regulation of intercellular adhesion and epithelial tubulogenesis.

First, we show that both Daam1 and E-cadherin localize to cell-cell contacts during nephron development and that Daam1 is required for promoting E-cadherin localization at sites of cell-cell contact. In contrast to what we observed during these early stages of nephron development following the depletion of Daam1, in mature nephrons, we were unable to detect junctional Daam1, and we likewise did not observe the effect on localization of E-cadherin. These data suggest that Daam1 is necessary for efficient localization of E-cadherin at cell junctions in nephron progenitor cells during early stages of nephron morphogenesis.

This conclusion builds upon certain assumptions. For example, it is possible that the overexpression of GFP-Daam1 has an impact on our localization analyses, or that cells depleted of Daam1 in mature nephrons are starting to recover due to a decrease in available morpholino pools with time. However, multiple lines of evidence suggest that both of these scenarios are highly unlikely. In the first case, prior studies indicate that the overexpression of full length Daam1 has little to no affect the actin cytoskeleton or other cellular processes (Liu et al., 2008). Additionally, several studies demonstrate that the distribution of the GFP-Daam1 recapitulates its endogenous localization as discerned via immunostaining (Jaiswal et al., 2013; Li et al., 2011; Nishimura et al., 2012, 2016). In the second case, consistent with our observations in *Xenopus* nephrons, sub-confluent cultures of MDCK cells stably expressing shDaam1 show transient repression of E-cadherin localization to cell-cell contacts that appears to be lost as the cells become confluent.

Indeed, our data support recent findings suggesting that actin nucleating proteins and Rho GTPases are required for early stages of E-cadherin mediated cell-cell adhesion in MDCK epithelial cells and not the maintenance of mature junctions (Collins et al., 2017). They also imply that morphological defects that we see in mature nephrons are consequences of earlier events. It is interesting to note that in similar fashion the silencing of E-cadherin expression in MDCK cells disrupts formation of cell-cell junctions whereas its signaling seems to be largely dispensable in already established epithelium as long as the cells are not mechanically stressed (Capaldo and Macara, 2007).

Second, we show that Daam1 localizes to actin protrusions and newly formed cell-cell contacts in developing nephron, suggesting that Daam1 via E-cadherin coordinates the assembly of cell-cell junctions during CE. Actin polymerization at the cell’s membrane mediates polarized movement and intercalation of cells during CE through engagement of cadherins (Huebner and Wallingford, 2018; Huebner et al., 2020). Here, we show that Daam1 activity functions to ensure proper organization and size of nephrogenic primordium at the time of CE as well as polarized movement of renal epithelial sheets. Our data support previous studies implicating Daam1 in CE and polarized cell movements (Ang et al., 2010; Kida et al., 2007; Liu et al., 2008). Furthermore, we show that Daam1 mediates polarized movement and cohesion of MDCK cells without slowing down the motility of individual cells. Taken together these data suggest that Daam1 promotes collective cell movements by controlling actin’s polymerization at cell-cell contacts and strengthening of E-cadherin-based adhesion.

Finally, Wnt9b and Wnt11 regulate tubular nephron morphogenesis, and disruptions in the cell behaviors traditionally regulated by the PCP pathway define their loss-of-function phenotypes (Karner et al., 2009; O’Brien et al., 2018). We show that inhibiting signaling activity of Daam1 in the prospective nephron results in a set of phenotypic characteristic comparable to those reported for Wnt11 and Wnt9b. Moreover, earlier studies have demonstrated that Daam1 can rescue Wnt11-induced CE gastrulation defects (Liu et al., 2008). These data collectively point to Daam1 as a potential downstream effector of Wnt11 and Wnt9b signaling in the control of nephric tubulogenesis. Interestingly, *wnt9b* and *wnt9a* are also induced in the injured nephrons and a mutation in *fzd9b* is associated with reduced regenerative capacity of nephric tubules (Kamei et al., 2019). These findings also suggest a potential role for Daam1 in repair and regeneration of nephric tubules. This hypothesis is further supported by increasing research evidence that the actin-based protrusions are important for the repair of cell-cell junctions (Li et al., 2020) and studies showing that Daam1 promotes assembly of actin-based protrusions (Jaiswal et al., 2013; Nishimura et al., 2016).

Cadherin localization is a complex process, and current studies propose at least three different ways of achieving efficient localization of E-cadherin: (1) clustering enforced by cortical F-actin, (2) clustering regulated by exposure of neighboring cells through their actin-based protrusions and (3) clustering promoted by condensation of F-actin networks via myosin (Yap et al., 2015). However, further research into timely regulation of E-cadherin is required to characterize the precise molecular details and specify relationships between different modes of E-cadherin clustering. Ultimately, investigations into how regulation of junctional E-cadherin by Daam1 fits in with different modes of E-clustering will also be important to examine in the future.

## ACKNOWLEGMENTS

We thank Miller lab members as well as Dr. Pierre McCrea and Dr. Jae-II Park and their lab members for lively discussions and suggestions on the manuscript. We also thank Dr. Richard Behringer, Dr. Yoshihiro Komatsu, Dr. Oleh Pochynyuk and Dr. Anna Marie Sokac, for their input and suggestions on the project. We thank Kenneth Dunner Jr. at the UT MD Anderson High Resolution Electron Microscopy Facility and CCSG grant NIH P30CA016672 for support with the transmission electron microscopy data. We are also thankful to Raymond Habas, Bruce Goode, Norihiro Sudou and Masanori Taira for providing antibodies and constructs. We thank the instructors and teaching assistants of the 2015 National *Xenopus* Resource Advanced Imaging Course. We are grateful to J.C. Whitney and T.H. Gomez who took care of the animals, even during Hurricane Harvey. We are grateful to the UTHealth Office of the Executive Vice President and Chief Academic Officer and the Department of Pediatrics Microscopy Core for funding the Zeiss LSM800 confocal microscope. We also thank BSRB Microscopy Facility at Department of Genetics, UT-MD Anderson Cancer Center. This work was funded by National Institute of Diabetes and Digestive and Kidney Diseases grants (K01DK092320, R03DK118771 and R01DK115655 to R.K.M.), startup funding from the Department of Pediatrics, Pediatric Research Center at the McGovern Medical School (to R.K.M.), The Antje Wuelfrath Gee and Harry Gee, Jr. Family Legacy Scholarship (to V.K.S.), and The Gigli Family Endowed Scholarships (to V.K.S.)

## AUTHOR CONTRIBUTIONS

V.K.S. conceived of the project, preformed experiments, analyzed data and wrote the original draft of the manuscript. M.E.C. cloned pCS2-mCherry-Daam1 and pCS2-mCherry-Daam1 (Ile698Ala) constructs and contributed to data validation. V.K.S., M.E.C., A.B.G. and R.K.M. preformed experiments to generate MDCK shDaam1 cells. V.K.S. and A.P. imaged the wound healing assays, FRAP experiments and analyzed FRAP data. V.K.S. and M.K. conducted TEM imaging analyses. A.B.G. and R.K.M. oversaw the experiments and supervised the project. All authors were involved in critical evaluation and editing of the manuscript.

## MATERIALS AND METHODS

### Xenopus laevis

*Xenopus laevis* adult male and female frogs were obtained from Nasco (LM00531MX and LM00713M) (Fort Atkins, WI, USA) and maintained according to standard procedures. *Xenopus* embryos were obtained by *in vitro* fertilization (Sive et al., 2000) and staged as previously described by Nieuwkoop and Faber (NF) (Nieuwkoop and Faber, 1994). All work was carried out in accordance with the University of Texas Health Science Center at Houston, Institutional Animal Care and Use Committee (IACUC) protocol #AWC-19-0081.

### MDCK cell lines

Madin-Darby Canine Kidney (MDCK) II cell lines were purchased from the American Type Culture Collection (ATCC). MDCK cells were cultured at 37°C with 5% CO2 in Dulbecco’s Modified Eagle’s Medium (DMEM) (Sigma, D6429) supplemented with 10% fetal bovine serum (FBS) (Sigma, F0926) and 1% Antibiotic-antimycotic solution (Sigma, A5955).

### Embryo microinjections

*Xenopus* embryos were microinjected at one-cell or into V2 blastomere at eight-cell stage, targeting embryonic kidney (DeLay et al., 2016; Moody and Kline, 1990; Nieuwkoop and Faber, 1994). Embryos were injected with synthetic mRNAs alone or in combination with antisense morpholino oligonucleotides (MOs). For mRNA injections, capped mRNA transcripts were synthesized from DNA-plasmids using SP6 mMessage mMachine transcription kit (ThermoFisher, AM1340M) and purified. pCS2-GFP-Daam1 and pCS2-GFP-Daam1 (Ile698Ala) plasmids were a gift from Dr. Raymond Habas’s and Dr. Bruce Goode’s labs, respectively (Lu et al., 2007). A mutation A2822G discovered in these plasmids was corrected by site directed mutagenesis as previously reported (Corkins et al., 2019) prior to mRNA synthesis. pCS2-membrane-tagged-RFP (mRFP)(Davidson et al., 2006), pCS2-membrane-tagged-EGFP (mEGFP)(Shindo and Wallingford, 2014) and pCS2-mCherry-Utrophin (mCherry-UtrCH)(Burkel et al., 2007) constructs were gifts from Dr. Raymond Keller’s lab, Dr. John Wallingford’s lab and Dr. William Bement’s lab, respectively. Formerly developed translation-blocking Daam1 (5′GCCGCAGGTCTGTCAGTTGCTTCTA 3′) (Corkins et al., 2018; Habas et al., 2001; Miller et al., 2011) and standard control (5′CCTCTTACCTCAGTTACAATTTATA 3′) MOs were purchased from GeneTools, LLC (Philomath, OR, USA). MOs were injected at 20ng per embryo while the amount of injected mRNA per embryo were as follows: GFP-Daam1 [1ng], mCherry-Daam1 [1ng], GFP-Daam1(I698A) [1ng], mCherry-Daam1(I698A) [1ng], mRFP [0.5ng], mGFP [0.5ng] and mCherry-UtrCH [1ng].

### Generation of stable MDCK shDaam1 cell lines

shDaam1 knockdown cell lines were generated by a retrovirus-based transduction method as described (Corkins et al., 2019). Briefly, the pLKO.1 lentiviral shDaam1 constructs (TTTCAGGAGATAGTATTGTGC, AAACAGGTCTTTAGCTTCTGC) were purchased from GE-Dharmacon (Clone ID: TRCN0000122999, Clone ID: TRCN0000123000). HEK293T cells were co-transfected with shDaam1 and virus packaging plasmids (psPAX2 and pMD2.G) using Polyethylenimine (PEI). The viral titers were collected starting 24 hours post transfection over the course of two days and purified using 0.22 μm Polyethersulfone (PES) syringe filters. Infections were carried out in the presence of polybrene. MDCK II cells remained in infection media for 24 hours, followed by puromycin selection with final concentration of 0.70 μg/ml. Lastly, MDCK II shDaam1 knockdown stable cell lines were validated by Western blotting.

### Western blotting

Western blotting was carried out using published protocols (Kim et al., 2002; Williams et al., 2017). In short, *Xenopus* embryos were cultured to desired stage and collected. The whole-embryo lysates were prepared by resuspending 10-20 embryos in a prechilled TX100-lysis buffer (10 mM HEPES, 150 mM NaCl, 2 mM EDTA, 2 mM EGTA, 0.5% Triton X-100, pH 7.4) and centrifuged at 18,407 RCF at 4°C for 5 minutes. The resulting protein lysates were resuspended in an equal volume 2X Laemmli (BioRad,161-0737) + dithiothreitol (Fisher BioReagents, BP17225) solution, and incubated at 95°C for 2 minutes. For making protein lysates using MDCK cells, cells were washed with PBS and pelleted by centrifugation at 18,407 RCF at 4°C for 5 minutes. Cell pellets were resuspended in in a prechilled Triton-lysis buffer (50 mM Tris pH 7.4, 1% Triton X-100, 150 mM NaCl, 1 mM EDTA, 1 mM EGTA, 1 mM PMSF, 1 Mm Na_3_VO_4_, 10 mM sodium fluoride, 10 mM β-glycerophosphate, 1 mg/ml aprotinin, and 1 mg/ml leupeptin). The cell lysates were incubated on ice for 20 minutes, followed by sonication and centrifugation at 18,407 RCF for 10 minutes at 4°C. Bradford assay was used to determine the total amount of protein in the lysates. The protein samples were run on an 8% SDS-PAGE gel and transferred to nitrocellulose or polyvinylidene difluoride (PVDF) membranes. The blots were blocked for at least 1.5 hours at the room temperature using the KPL Detector Block Kit (Sera Care, 5920-0004, 71-83-00) and probed with primary antibodies overnight at 4°C. The next day, after series of washes with Tris-buffered saline containing 0.1% Tween 20 (TBST), blots were incubated with secondary antibodies for at least 1 hour at the room temperature. Protein expression levels were detected with enhanced chemiluminesence (SuperSignal West Pico PLUS Chemiluminescent Substrate, Thermo Fisher, 34580) using LiCor and BioRad ChemiDoc XRS imagers. The following antibodies were used: rabbit anti-Daam1 (1:1000, Proteintech, 14876-1-AP), rabbit anti-Daam1 (1:1000, gift from Dr. Raymond Habas), rabbit anti-GAPDH (1:1000, Santa Cruz, sc-25778), mouse anti-E-cadherin (1:1000, BD Transduction Laboratories, 610182), rabbit anti-GFP (1:250, ICL Lab, RGFP-45A), anti-rabbit IgG (H + L)-HRP (1:5000, BioRad,1706516) and anti-mouse IgG (H + L)-HRP (1:5000, BioRad, 1706516).

### Immunostaining and staining

*Xenopus* embryos were fixed in MEMFA (3.7% formaldehyde, 4 mM MOPS, 2 mM EGTA, and 1 mM MgSO_4_, pH 7.4) for 1 hour at room temperature or overnight at 4°C. Embryos to be stained with Phalloidin, were fixed using methanol-free formaldehyde (Thermo Scientific, 28908). Immunostaining was carried out according to previously published methods (Hemmati-Brivanlou and Melton, 1994; Krneta-Stankic et al., 2010). In short, fixed-embryos were washed 3 times for 15 minutes at room temperature with phosphate buffered saline (PBS) containing 0.1% Triton X-100 and 0.2% bovine serum albumin (BSA) and blocked using 10% goat serum diluted in PBST for 1 hour at room temperature. Embryos were incubated with the primary antibodies overnight at 4°C. The next day, embryos were washed 5 times for 1 hour with PBS-T at room temperature and then incubated with the secondary antibodies overnight at 4°C. The following day, embryos were washed 3 times for 1hr at room temperature and dehydrated in methanol prior to clearing. To preserve Phalloidin-labeling, isopropanol was used to dehydrate Phalloidin-stained embryos (Nworu et al., 2014). Embryos were cleared using BABB (1-part benzyl alcohol: 2-parts benzyl benzoate) clearing solution and imaged. MDCK cells were fixed in 4% paraformaldehyde (PFA) for 10 minutes at room temperature. Cultures were washed 3 times with PBS and incubated with 50mM ammonium chloride for 10 minutes at room temperature to neutralize the PFA. Next, samples were washed 3 times with PBS and blocked using 10% goat serum/ PBST for 1 hour and incubated with primary antibodies overnight at 4°C. The following day, MDCK cells were washed 3 times with PBS and incubated with secondary antibodies for 1 hour at room temperature. Lastly, stained samples were washed again 3 times with PBS prior to mounting in Fluoromount-G medium (Southern Biotech, 0100-01) for imaging. The following primary antibodies were used: chicken anti-GFP (1:250, Abcam, ab13970), rabbit anti-RFP (1:250, MBL International, PM005), rabbit anti-GFP (1:250, ICL Lab, RGFP-45A), rabbit anti-Lhx1 (1:250, gift from Dr. Masanori Taira) and mouse anti-E-cadherin (1:100, BD Transduction Laboratories, 610182). For detection of primary antibodies the following secondary antibodies were used: anti-rabbit IgG Alexa 488 (1:500, Invitrogen, A-11008), anti-rabbit IgG Alexa 555 (1:500, Invitrogen, A-21428), anti-rabbit IgG Alexa 647 (1:500, Invitrogen, A-21244), anti-mouse IgG Alexa 488 (1:500, Invitrogen, A-11001), anti-mouse IgG Alexa 555 (1:500, Invitrogen, A-21422), anti-mouse IgG Alexa 647 (1:500, Invitrogen, A-21235), anti-mouse IgG Alexa 488 (1:500, Jackson ImmunoResearch, 715-545-150) and anti-chicken IgY Alexa 488 (1:500, Invitrogen, A-11039). Fluorescent probes used for staining were as follows: FITC-conjugated lectin from *Erythrina cristagalli* (1:500, Vector labs, FL-1141), Phalloidin-Alexa 568 (embryos-1:40 and cells-1:200, Invitrogen, A12380), and diamidino-2-phenylindole (DAPI) (1:500, Thermo Scientific, 62247).

### Transmission electron microscopy (TEM)

*Xenopus* embryos were fixed in 2%formaldehyde+0.5% glutaraldehyde (Ted Pella Inc., 18505 and 18426). Fixed embryos were washed with 0.1 M sodium cacodylate buffer and treated with 0.1% Millipore-filtered cacodylate buffered tannic acid. Embryos were post-fixed using 1% buffered osmium, followed by staining using 1% Millipore-filtered uranyl acetate. Stained embryos were dehydrated by washing in increasing concentrations of ethanol, permeated, and embedded in LX-112 medium. After embedding, embryos were placed in a 60°C oven for approximately 3 days to polymerize. Polymerized samples were sectioned using Leica Ultracut microtome (Leica, Deerfield, IL). Collected ultrathin sections were stained with uranyl acetate and lead citrate in a Leica EM Stainer and subjected to imaging.

### Isolation and live imaging of *Xenopus* kidney cells

Embryos were microinjected with 1ng GFP-Daam1 mRNA into the V2 blastomere, targeting kidney. Around embryonic NF stage 30, GFP-positive nephrons were surgically dissected under a fluorescent stereomicroscope using a pair of sharpened forceps (Fisher, NC9404145). Microsurgical dissections were performed in plastic petri dishes coated with 2% agar containing Danilchik’s for Amy (DFA) solution (53 mM NaCl, 5 mM Na_2_CO_3_, 4.5 mM Potassium Gluconate, 32 mM Sodium Gluconate, 1 mM CaCl_2_, 1 mM MgSO_4_, buffered to pH 8.3 with 1 M bicine) supplemented with 1 g/l of and Antibiotic antimycotic solution (1:100, Sigma, A5955). To dissociate into single cells, the isolated nephrons were transferred to fibronectin coated (1μg/ml, Roche, 10-838-039-001) glass-bottom imaging chambers (Thermo, A7816) prefilled with Calcium Magnesium Free Media (CMFM) (88 mM NaCl, 1 mM KCl, 2.4 mM NaHCO_3_, 7.5 mM Tris pH 7.6). After ∼30 minutes at the room temperature, as much as possible of the CMFM media was aspirated from top of the chamber without disturbing the cells. Fresh DFA media was added and again removed by careful aspiration. In order to ensure complete removal of the CMFM media, this process was repeated at least 5 times. The cells were left undisturbed at the room temperature for 15-30 minutes prior to imaging.

### Wound-healing assay

MDCK cells were seeded in 6-well culture plates at a density of 100,000 cells/well and allowed to reach confluency. Prior to inducing a “wound” or scratch or in a confluent cell monolayer, the cells were treated with Mitomycin C (10 μg/ml) for 3 hours at 37°C to prevent future proliferation. A linear scratch was made by gliding the 200 μl sterile tip across the bottom of each well. After making a scratch, cells were washed with 1X PBS and refed with 5 ml of 10%FBS/DEMEM supplemented with 1%Antibiotic-antimycotic solution. Samples were placed in 37°C heated imaging chamber with 5% CO2 and subjected to time-lapse imaging.

### Fluorescence Recovery after Photobleaching (FRAP)

To visualize actin dynamics, embryos were injected into V2 blastomere at 8-cell stage with 1 ng mCherry-UtrCH mRNA along with 20 ng of Daam1 or Standard (control) MO. Injected embryos were cultured to around NF stage 30. Since the opaqueness of *Xenopus* epithelium prevents direct *in vivo* imaging of developing nephron at this early embryonic stage, “windowed” embryos were generated. To make windowed embryos, embryos were anesthetized in 0.04% (0.15%) Ethyl 3-aminobenzoate methanesulfonate (Sigma, E10521) diluted in DFA. Next, the epithelium covering developing nephron was surgically removed under fluorescence-dissecting microscope, exposing developing nephron for *in vivo* imaging. Windowed embryos were mounted under glass-cover slips and subjected to FRAP. FRAP assays were performed on an inverted 3i spinning disk microscope integrated to a NIKON TiE with perfect focus and equipped with Vector™ FRAP scanning module and a Hamamatsu Flash 4.0 camera. Images were acquired with a NIKON Plan-Apo 60X water 1.2 NA objective. A small fragment at the midpoint of a cell-cell junction was bleached with the 561nm laser line at 100% power. A series of twenty pre-bleach images were captured and post-bleach recovery was recorded continuously until fluorescent signal reached a steady state. Movies were analyzed using Slidebook 6.0 and curve fitting was done with SigmaPlot and GraphPad Prism 8.0 software. For curve fitting, a single exponential function (f(t)=α(1-e^-kT^), where T^1/2^ (half-time of recovery) is ln 0.5/(-k), and α is the mobile fraction was used. Raw recovery curves were corrected for background and photofading. Lowest fluorescence signal and the time-point after bleaching were scaled to 0 and curves were normalized to 1 based on the reference signal before bleaching. Per embryo, between 1 and 5 single junctions were photobleached. To avoid bleaching-induced variation in fluorescence, junctions picked for photobleaching were spaced far apart. All FRAP experiments have been carried out using multiple embryos and repeated at least three times.

### Image acquisition and processing

Olympus SZX16 fluorescent stereomicroscope equipped with Olympus DP71 camera was used for carrying out *Xenopus* microsurgical manipulations, embryo mounts and scoring of kidney phenotypes in NF stage 40 embryos. Zeiss LSM800 microscope with Airyscan detector, LeicaSP5 and NikonA1 were used for confocal imaging of fixed and live samples. Captured images and time-lapse movies were exported as original files and processed using ImageJ (Fiji plugin). Image panels were built using FigureJ plugin. Final figures were assembled in Adobe Photoshop CC. TEM imaging was carried out in a JEM 1010 transmission electron microscope (JEOL, USA, Inc., Peabody, MA) with the AMT Imaging System (Advanced Microscopy Techniques Corp, Danvers, MA).

### Quantification and statistical analyses

ImageJ (Fiji plugin) software was used for quantitative image analyses. For quantification of E-cadherin and F-actin staining **(Figures 3, 5 and 6)** images were captured using the same settings. The mean fluorescence intensity along the length of selected junction was measured. To analyze tissue organization during nephron development **(Figure 3)** cells were manually counted using the Cell Counter tool. The shortest distance between two neighboring nuclei was measured using the straight-line selection tool. The area and circularity were measured using the Analyze Particles tool. Wound area **(Figure 7)** was also measured using the Analyze Particles tool. To obtain traveled distance, trajectories of randomly selected cells were traced manually in each frame of the time-lapse using Plugin for Motion Tracking and Analysis (MTrackJ) (The Biomedical Imaging Group of Erasmus University Medical Centre, Rotterdam, The Netherlands). All experiments were repeated at least two times, with the exception of TEM studies, which due to the prohibitive costs of the experiment represent single-trial analyses. The exact sample size and statistical analysis for each experiment are presented in the corresponding figure legend. Statistical analyses were carried out using GraphPad Prism 8.0 software.

