## Supplementary material for "Cell-Cell Adhesion During Nephron Development Is Driven by Wnt/PCP Formin Daam1": figures supplemental

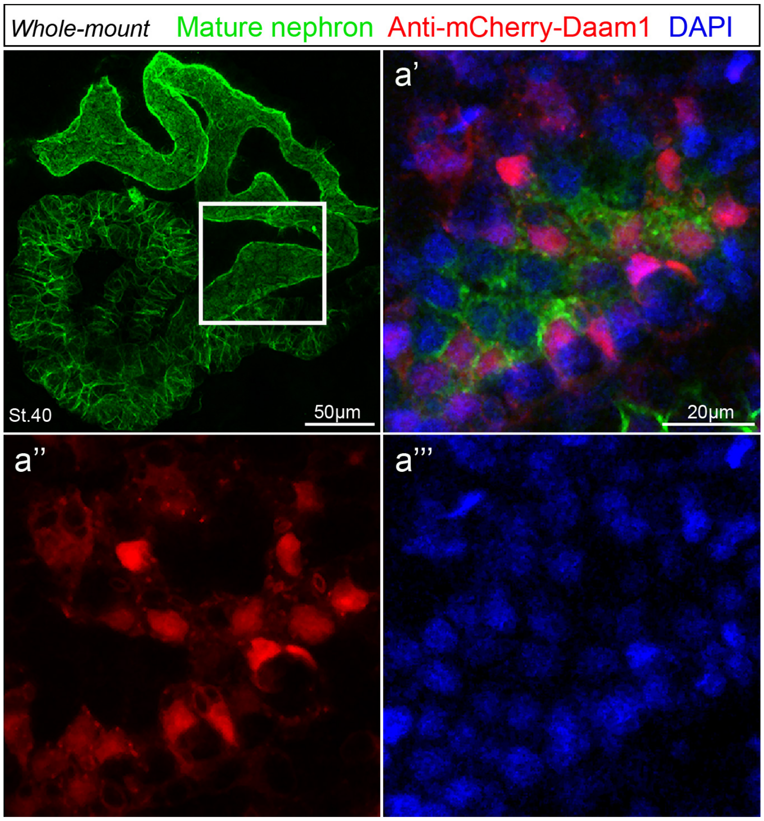


**Figure S1. Daam1 is absent from cell-cell junctions in fully developed embryonic nephron**

Whole-mount immunostaining of mature *Xenopus* embryonic nephron labeled with 3G8 and 4A6 (green), mCherry to visualize Daam1(red) and DAPI staining labeling nuclei (blue). a’-a’’’ close-up images of white box.

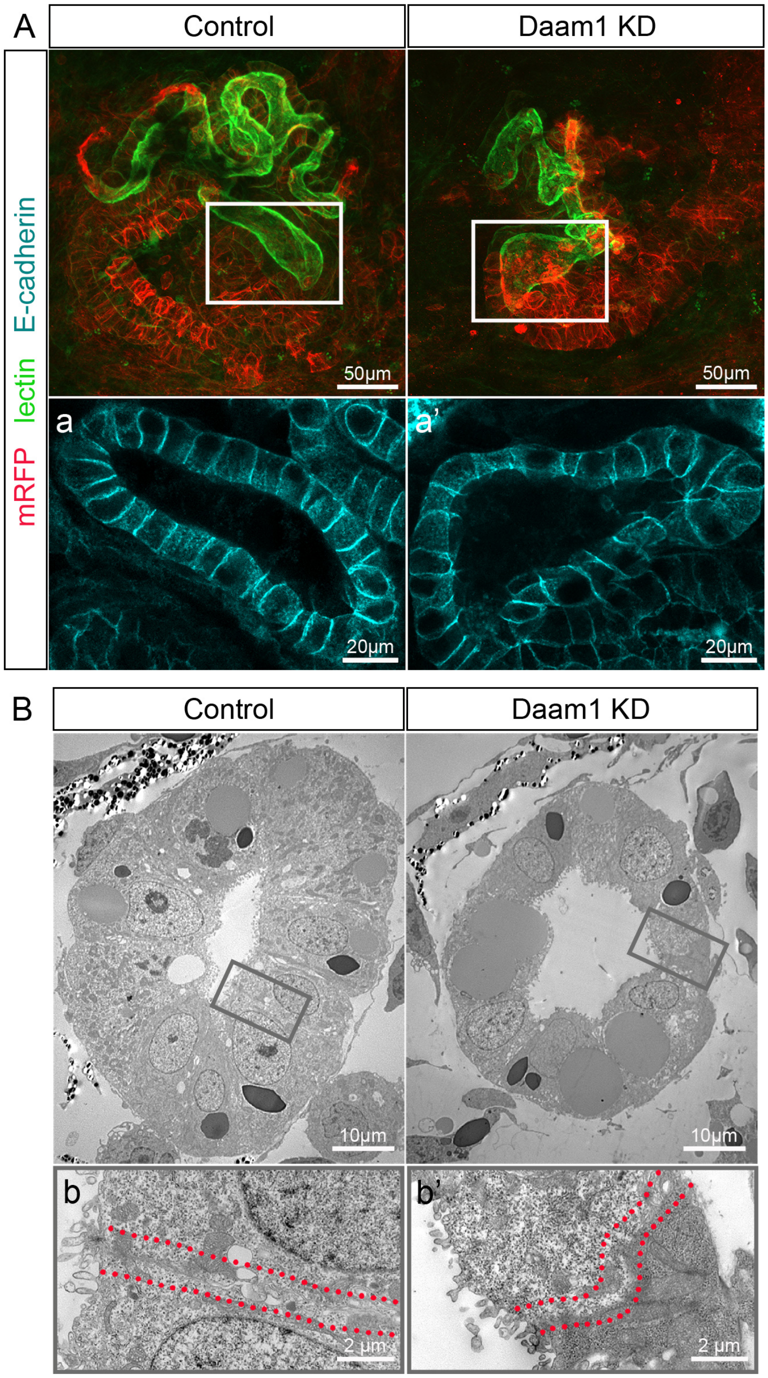


**Figure S2. Loss of Daam1 results in cell shape and tissue architecture abnormalities in mature nephrons**
Kidney-targeted morpholino microinjections were carried out to manipulate the expression levels of Daam1. Control (Standard) or Daam1 antisense morpholinos were co-injected with membrane tagged RFP (mRFP) mRNA as a linage tracer. Analyses of mature nephrons by confocal and Transmission Electron Microscope (TEM) imaging show that decrease in Daam1 signaling levels affects the size and shape of nephric cells.
(A) Maximum projection confocal images of mature nephrons labeled with lectin (green) and antibodies against mRFP (red) and E-cadherin (cyan). a-a' - close-up images corresponding to regions in the white boxes, see Videos S5 and S6.
(B) TEM images show cross-section samples of the Control and Daam1-depleted nephrons. b-b' - close-up images corresponding to regions in the gray boxes. The red dotted lines outline morphology of intercellular junction.

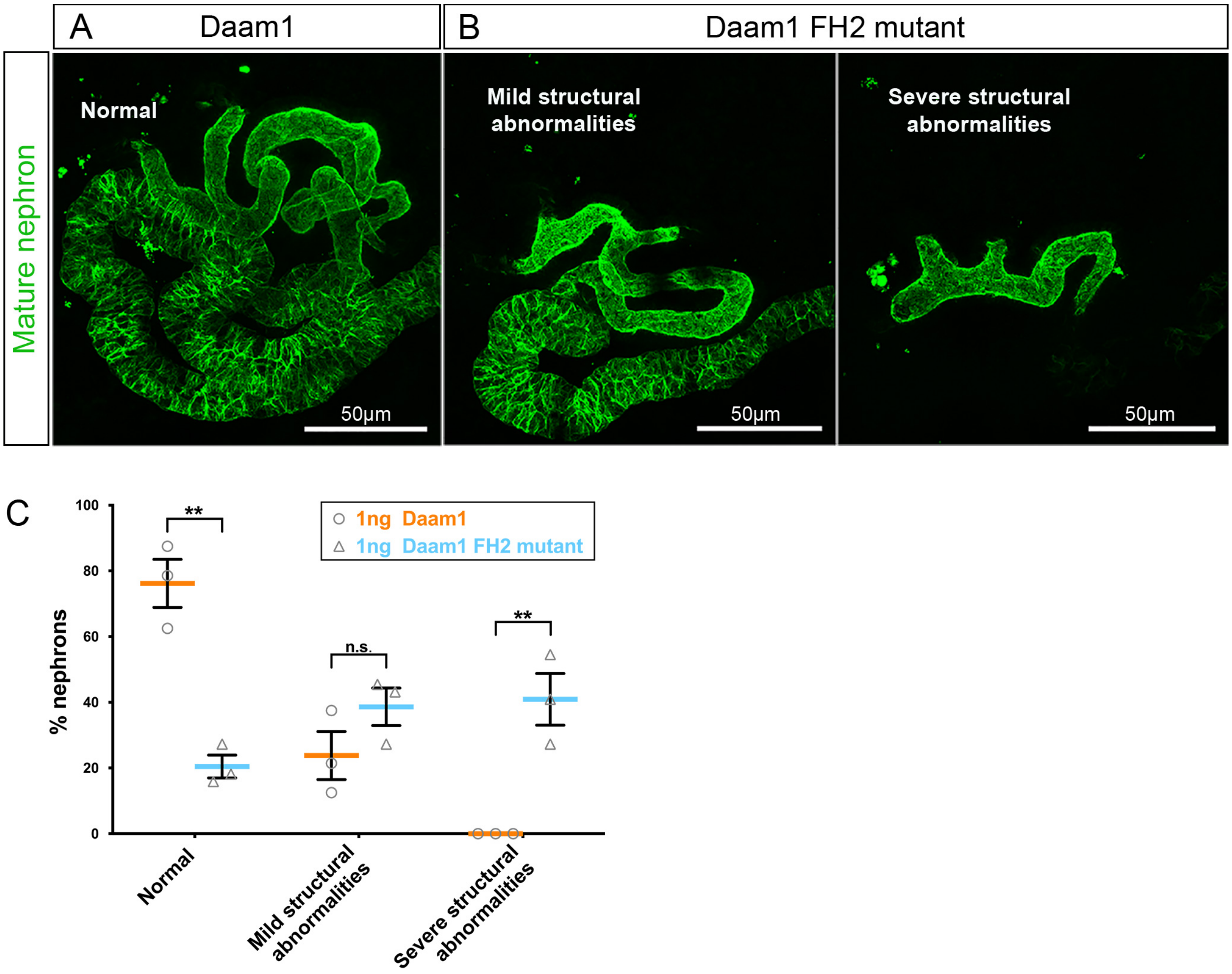


**Figure S3. FH2 domain of Daam1 mediates nephron morphology**

Nephric progenitors expressing the full length Daam1 mRNA ultimately develop structurally normal nephrons in comparison to nephric progenitors expressing Daam1 FH2 mutant mRNA. Maximum projection confocal images of mature nephrons (NF stage 39-40) visualized by 3G8 and 4A6 antibodies (green) in embryos injected with 1ng Daam1 or 1ng Daam1FH2 mutant mRNA. Embryo microinjections were carried out at the 8 cell-stage into V2 blastomere fate-mapped to pronephric primordium.

(A) Representative image of mature nephron from embryos injected with 1ng Daam1 mRNA.

(B) Representative images of mature nephrons from embryos injected with 1ng Daam1 FH2 mutant mRNA.

(C) The graph represents quantification of the phenotypic severity of mature nephrons in embryos from A and B. $N_{Daam1}$=54 embryos across 3 experiments and $N_{Daam1FH2mutant}$ = 66 embryos across 3 experiments. Error bars indicate S.E. of the mean. $^{n.s.}$P=0.1234, $^{*}$P<0.0332, $^{**}$P<0.0021, $^{***}$P<0.0002 and $^{****}$P<0.0001 analyzed by unpaired t-test.

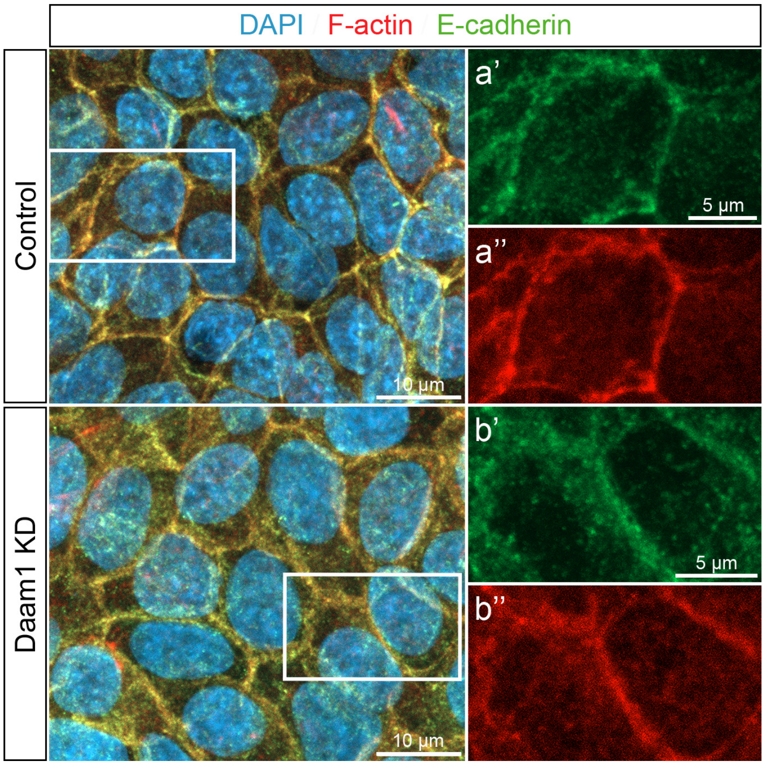


**Figure S4. E-cadherin localizes to shDaam1 depleted mature cell-cell junctions**
E-cadherin (green), F-actin (red) and DAPI (blue) in confluent the MDCK Control and shDaam1 knockdown cells. a'-a’’’ and b’-b’’’ represent close-up images of corresponding white boxes.

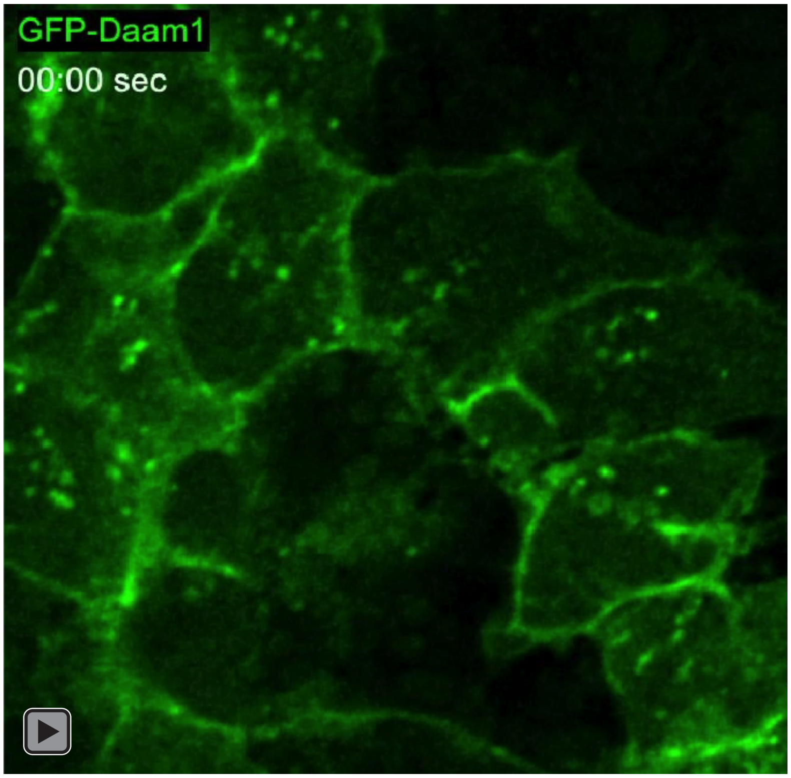


**Video S1.** Time-lapse of the nephric primordium expressing GFP-Daam1.

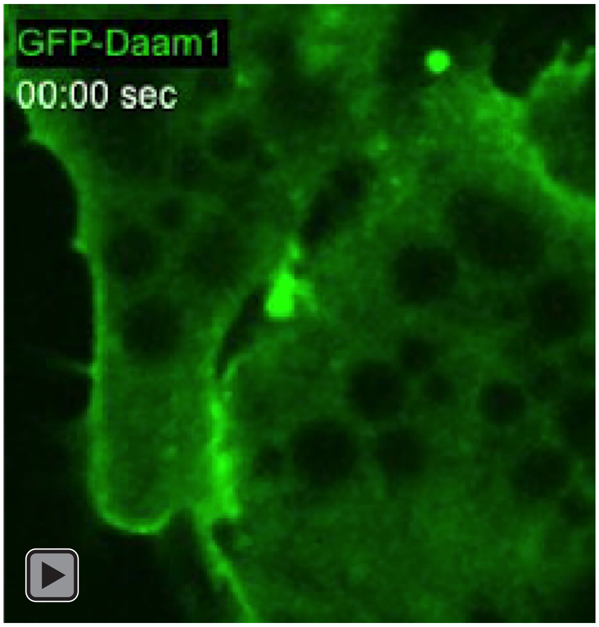


**Video S2.** Imaging of GFP-Daam1 localization during de-novo contact formation in cells isolated from the neprhic primordium expressing GFP-Daam1.

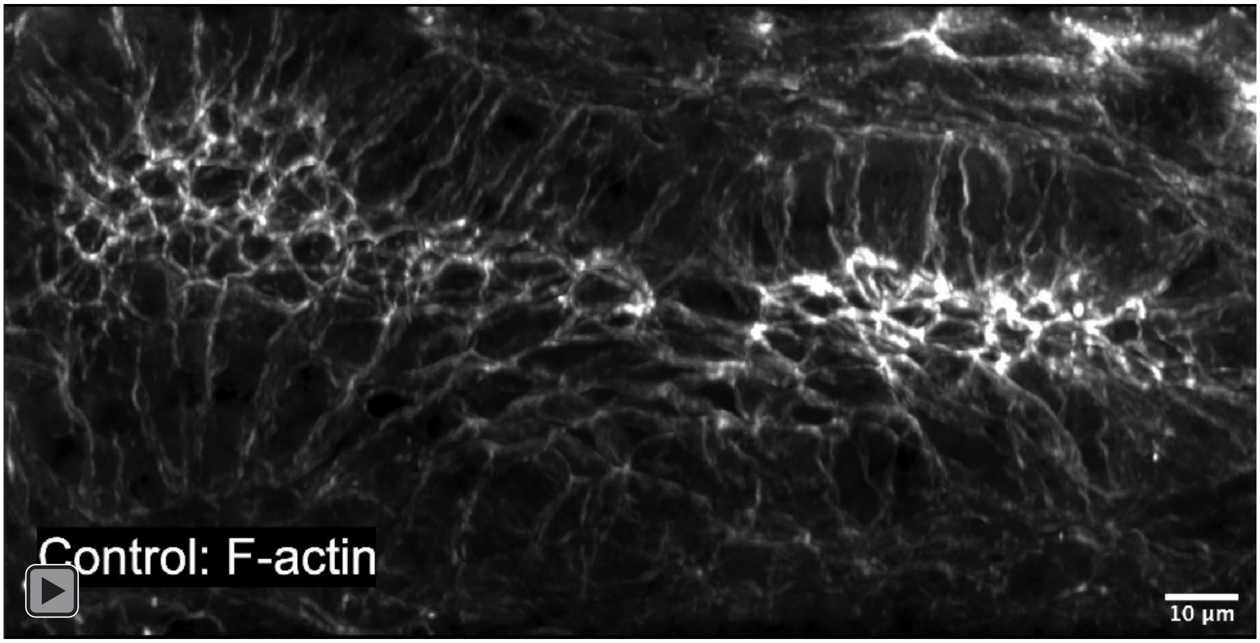


**Video S3.** 3D projection of sections from nephric primordium of Control embryos.

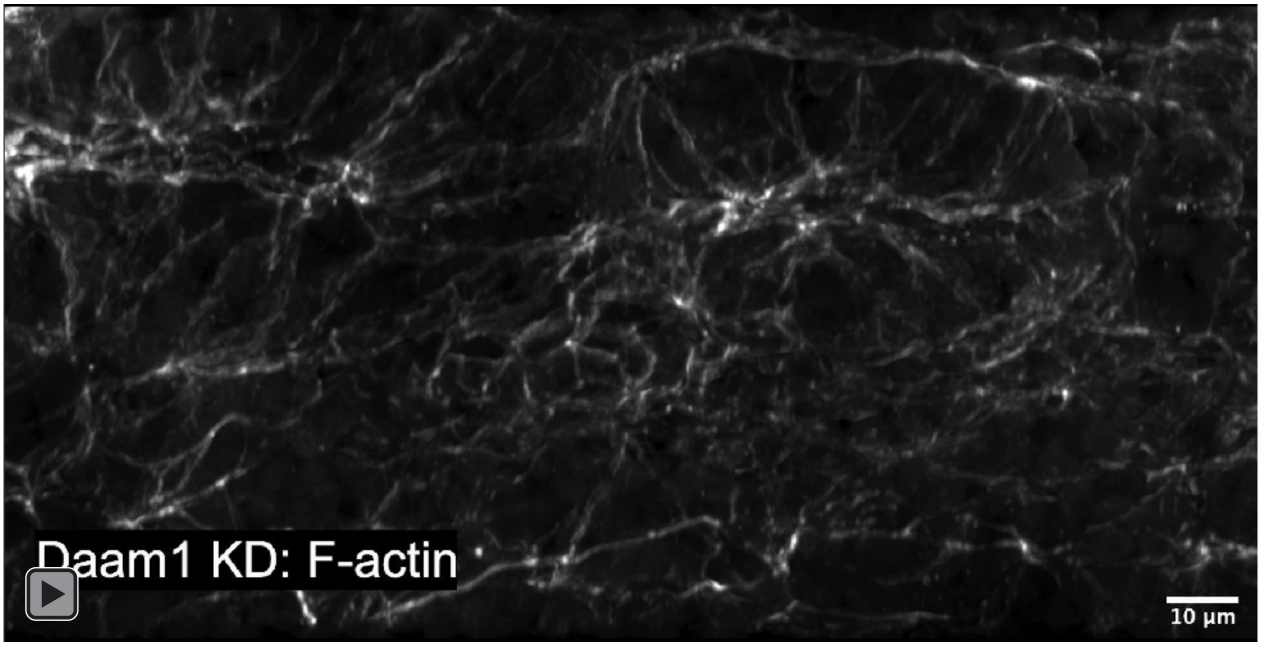


**Video S4.** 3D projection of sections from nephric primordium of Daam1 knockdown embryos.

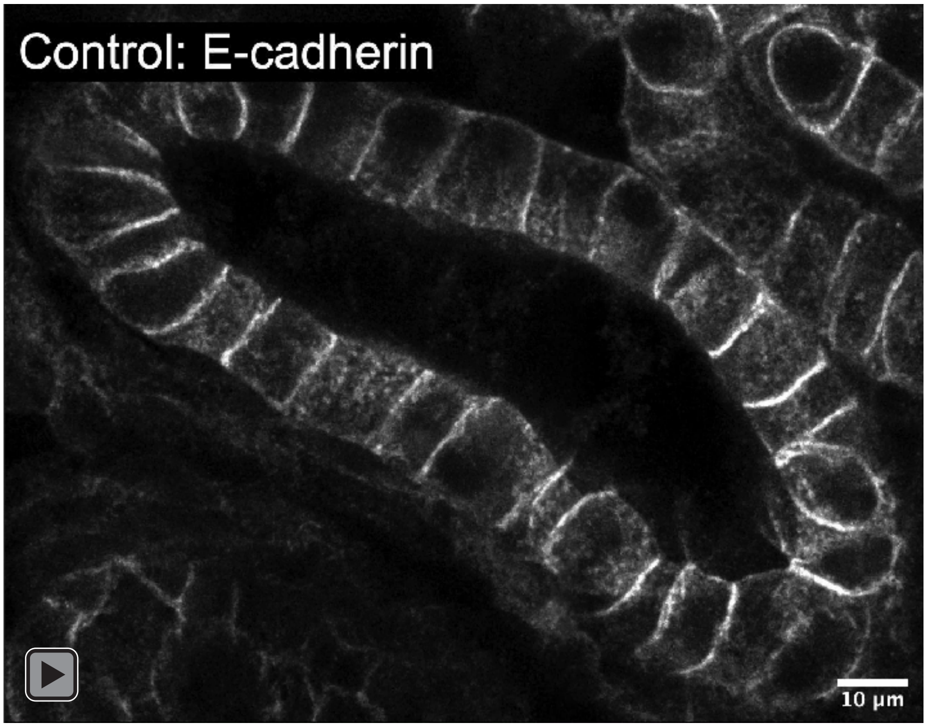


**Video S5.** 3D projection of sections from the mature Control nephron stained for E-cadherin.

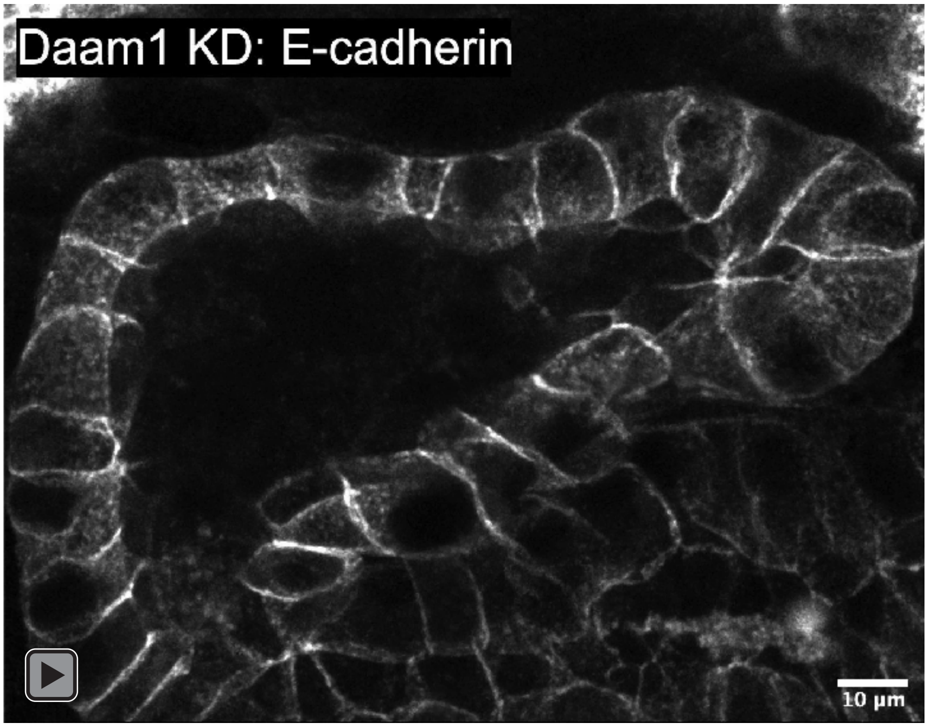


**Video S6.** 3D projection of sections from the mature Daam1-depleted nephron stained for E-cadherin.

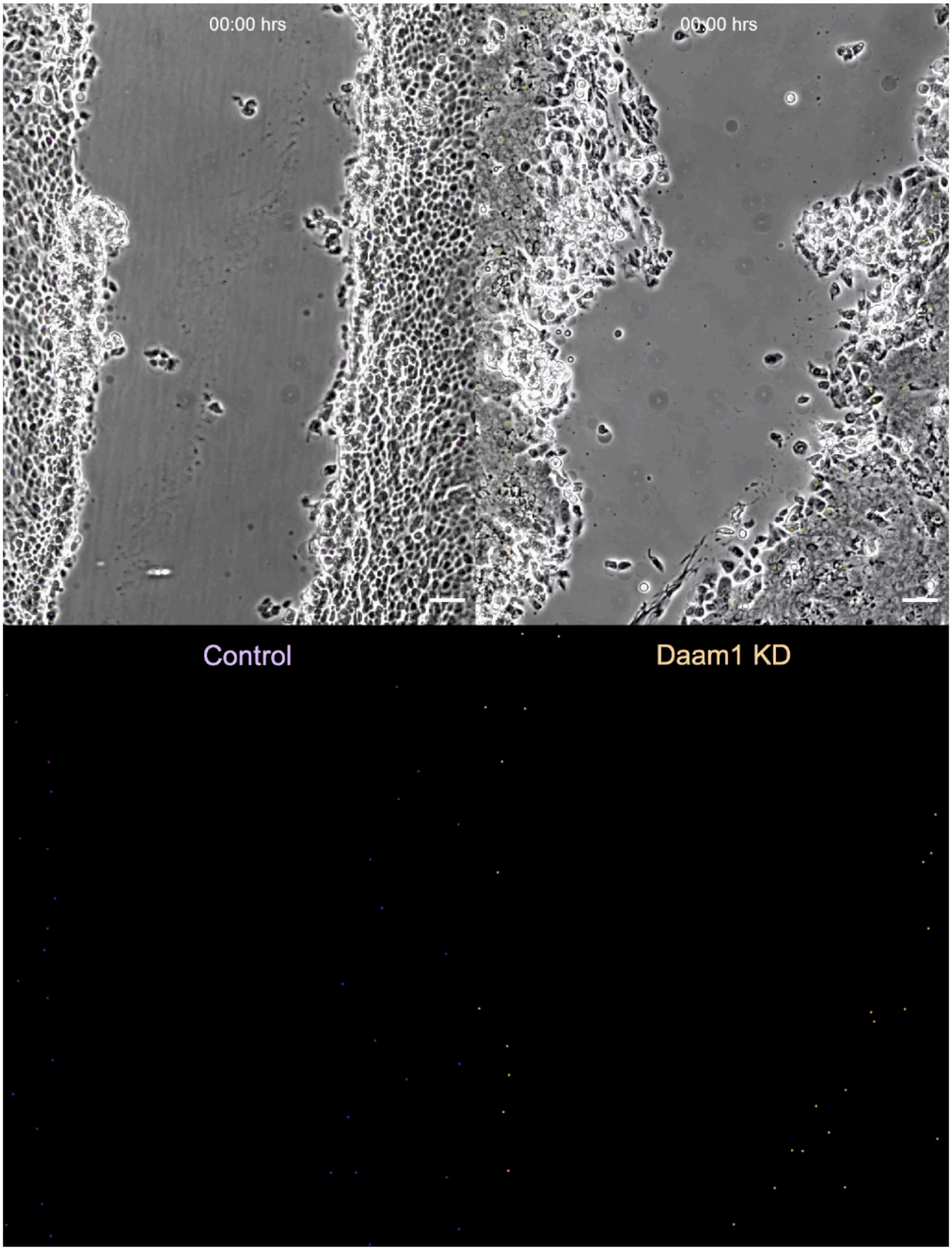


**Video S7.** Time-lapse of wound healing assay in the MDCK Control (left) and Daam1 KD (right) cells. Superimposed tracks show movement of individual cells during wound healing. Daam1 deficient cells exhibit a delay in a wound closure, uncoordinated movement and random detachment from migrating epithelial sheets in comparison with Control. Elapsed time is shown in hours at the top of the upper panels. Scale bars equal 50 microns.
